# Analysis of contributory gut microbiota and lauric acid against necrotic enteritis in *Clostridium perfringens* and *Eimeria* side-by-side challenge model

**DOI:** 10.1101/434449

**Authors:** Wen-Yuan Yang, Yuejia Lee, Hsinyi Lu, Chung-Hsi Chou, Chinling Wang

## Abstract

Gut microbiota has been demonstrated to be involved in intestinal nutrition, defense, and immunity, as well as participating in disease progression. This study was to investigate gut microbiota changes in chickens challenged with *netB*-positive *Clostridium perfringens* strain 1 (CP1) and/or the predisposing *Eimeria* species (*Eimeria*). In addition, the effects of lauric acid, a medium-chain fatty acid (MCFA), on NE reduction and modulation of microbiota were evaluated. The results demonstrated that microbial communities in the jejunum were distinct from those in the cecum, and the microbial community change was more significant in jejunum. Challenge of CP1 in conjunction with *Eimeria* significantly reduced species diversity in jejunal microbiota, but cecal microbiota remained stable. In the jejunum, CP1 challenge increased the abundance of the genera of *Clostridium sensu stricto 1*, *Escherichia Shigella*, and *Weissella*, but significantly decreased the population of *Lactobacillus*. *Eimeria* infection on its own was unable to promote NE, demonstrating decrements of *Clostridium sensu stricto 1* and *Lactobacillus*. Co-infection with CP1 and *Eimeria* reproduced the majority of NE lesions with significant increment of C*lostridium sensu stricto 1* and reduction in *Lactobacillus*. The changes of these two taxa increased the severity of NE lesions. Further analyses of metagenomeSeq, STAMP, and LEfSe showed significant overgrowth of *Clostridium sensu stricto 1* was associated with NE and *Eimeria* infection than *C. perfringens* challenge alone. The supplementation of lauric acid did not reduce NE incidence and severity but decreased the relative abundance of *Escherichia Shigella*. In conclusion, significant overgrowth of *Clostridium sensu stricto 1* in the jejunm is the major microbiota contributory to NE. Controlling proliferation of this taxon in the jejunum should be the niche for developing effective strategies against NE.

## Introduction

Necrotic enteritis (NE) as the result of proliferations of *Clostridium perfringens* (*C. perfringens*) type A and their associated toxins in the small intestine of chickens is a devastating enteric disease, characterized by sudden diarrhea, unexpected mortality, and mucosal necrosis [1, 2]. Up to 37% of commercial broiler flocks is estimated to be affected by this disease, and it has contributed to the losses of 6 billion dollars in the global poultry industry [3, 4]. In recent decades, *C. perfringens*-associated NE in poultry has been well-controlled by in-feed antimicrobial growth promoters (AGPs) [5]. However, the emergence of antibiotic-resistant bacteria from animals and the potential threat of transmission to humans has led to bans on using AGPs in many countries [6, 7]. Following the withdrawal of AGPs from poultry feed, NE has re-emerged as a significant disease to the poultry industry [8–11].

Gut microbiota is one of the central defense components in the gastrointestinal tract against enteric pathogens, which works by modulating host responses to limit the colonization of pathogens [12]. Interactions between gut microbiota and the host could influence intestinal morphology, physiology, and immunity [13]. Recently, gut microbiota has been demonstrated to regulate intestinal gene expression [14] and T cell-mediated immunity [15] as well as to accelerate the maturation of the gut immune system [16]. Conversely, a growing number of studies have observed gut microbial shifts in enteric diseases, considering that gut microbiota plays a role in the progress of disease development. Similar results were also represented in NE induction models, proposing that the disturbance of gut microbiota interacts with the host, subsequently promoting the development of NE [17–20]. In the case of human necrotizing enterocolitis, an enteric disease in infants associated with *C. perfringens* [21], a recent study found that *Bacteroides dorei*, an opportunist pathogenic bacterium in anaerobic infections, was associated with an increased mortality of this disease [22]. Furthermore, several studies have demonstrated that the increment of bacteria belong to a genus of *Escherichia-Shigella* was associated with *C. perfringens* infection [5, 20]. This evidence raised the possibility that certain microbes or microbiota in the gut may contribute to the virulence or development of enteric disease in chickens, particularly for NE.

The removal of AGPs drove the poultry industry to search for an alternative in prevention to decrease the incidence of NE. Probiotics, prebiotics, organic acids, plant extracts, essential oils, and enzymes arose in response to this demand, but the efficacy of those on NE reduction were variable and inconsistent [23, 24]. However, a medium-chain fatty acid (MCFA), lauric acid, was found to have strong *in vitro* antimicrobial activity against gram-positive organisms [25–27] and *C. perfringens* [28, 29]. In an *in vivo* trial, lauric acid with butyric acid demonstrated the lowest incidence and severity of NE compared to other treatments [29]. However, this promising result did not promote more applications of lauric acid against NE, and the interaction of lauric acid with gut microbiota was not even addressed. The evaluation of its modulation effect on NE reduction and gut microbiota simultaneously would be valuable in exploring specific microbial community contributory to NE.

Although *C. perfringens* is the causative etiological agent of NE, it is evident that other predisposing factors are required for NE induction [10, 30–32]. Even though gut microbiota has been suggested to be involved in the progress of NE development [33, 34], association between microbiota profile and NE development have not been well elucidated. Most studies intensively focused on changes of microbial communities in the ileum or cecum where higher quantity of microbes or/and more diverse microbial compositions were harbored; however, most results were inconclusive [17–19, 35, 36]. Inversely, microbiota in the jejunum, which serve as the primary site for colonization of *C. perfringens* and development of NE [33], was seldom evaluated. In the present study, we investigated gut microbiota targeting NE cases and in chickens with side-by-side treatments with the causative pathogen and parasitic predisposing factor, *Eimeria*, and expected to unveil the contributory microbe or microbiota to NE. The effects of lauric acid on NE reduction and modulation of microbiota were also examined to confer the alternative intervention to prevent and control NE.

## Materials and methods

### Ethics statement

All procedures for the care, housing and treatment of chickens were approved by the Institutional Animal Care and Use Committee at Mississippi State University (IACUC 16-439).

### Chicken, diet, and experimental design

A total of 50 male and female one-day-old unvaccinated broiler chicks (Cobb strain) were obtained from a commercial hatchery. Chicks were inspected on receiving to ensure their healthy status and randomly allotted to 5 groups, studying the NE incidence and gut microbiota after challenge of *netB*-positive *C. perfringens* (A group: CP1), co-infection with *netB*-positive *C. perfringens* and multi-species *Eimeria* (B group: CP1+*Eimeria*), addition of lauric acid to feed chickens co-infected with *netB*-positive *C. perfringens* and multi-species *Eimeria* (C group: CP1+*Eimeria*+LA), inoculation of multi-species *Eimeria* (D group: *Eimeria*), and no treatment (E group: CTL).

Chicks in groups were placed in separate temperature-controlled iron tanks with nets in the floor-pen facility and lined with fresh litter. Throughout the 19-day study period, wheat-based diets prepared based on the formula by Branton et al. [37] were offered for the first 7 days, and then the rations were replaced by fishmeal diets (wheat-based diets containing 50% fishmeal) obtained from the 1:1 mixture of wheat-based diet with fishmeal 60 N (Seven Springs Farm, Check, Virginia, USA), containing minimal 60% crude protein from days 8 until the end of the study. For the lauric acid supplementing group, 400 mg of lauric acid powder (Fisher Scientific, Pittsburgh, Pennsylvania, USA) was added into 1 kg of wheat-based diet or fishmeal diet to form the final ration for chickens from day 8 onward [29].

Co-infection with *netB*-positive *C. perfringens* (CP1) and multi-species *Eimeria* was applied to induce NE according to our previous studies. The success of reproducing NE was determined by clinical signs and intestinal lesion scores reaching 2 or more. In brief, chickens in the co-infection group were given a single gavage of coccidial inoculum at day 10, followed by oral administration of 3 ml CP1 inoculum with average 2.5×10^8^ colony-forming units (CFU)/ml at day 15 for 4 consecutive days with a frequency of 3 times daily. For a single challenge of CP1 or *Eimeria*, the same methodology and time points were conducted as in the co-infection group. All chickens were inspected on a daily basis and humanely euthanized at day 19 by carbon dioxide. Dead chickens not resulting from NE were excluded from the trial after necropsy. The experimental trial was reviewed and approved by the Mississippi State University Institutional Animal Care and Use Committee.

### Challenge strain and inoculum preparation

Anticoccidial live vaccine containing live oocysts of *E. acervulina*, *E. maxima*, *E. maxima MFP*, *E. mivati*, and *E. tenella* was used as a disposing factor. .The vaccine bottle contained 10,000 doses of oocysts in an unspecified proportion of *Eimeria* species. A ten-fold dose of vaccine was prepared then applied on *Eimeria*-treated and co-infection groups. *C. perfringens*, a clinical NE strain designated as CP1 obtained from Dr. John F. Prescott (Ontario Agricultural College, University of Guelph, Canada), was used to challenge chickens. This strain was characterized as *netB*-positive Type A and used to reproduce NE in a number of experiments [38–41]. CP1 was cultured on blood agar plates and incubated anaerobically at 37°C for overnight. A single colony was in turn transferred into 3 ml of fluid thioglycollate (FTG) medium (Himedia, Mumbai, Maharashtra, India) at 37°C for overnight. Thereafter, the bacterial suspension was inoculated into fresh FTG broth at a ratio of 1:10 and incubated at 37°C for 15, 19, and 23 hours, respectively. The whole broth cultures were used to induce NE based on the evidence that clostridia with toxins produce more severe disease than using cells alone [38]. The bacterial concentration (CFU/ml) of inoculum was calculated by plate counting using Brain Heart Infusion agar (Sigma-Aldrich, St. Louis, Missouri, USA), followed by anaerobic incubation at 37°C for 16 hours.

### Sample collection and lesion scoring

Three chickens per group were randomly selected to collect fecal contents from the jejunum (AJ, BJ, CJ, DJ, and EJ) and cecum (AC, BC, CC, DC, and EC). Among three CP1-challenged groups (A, B, and C), chickens suffering NE (lesion score ≥ 2) were preferentially collected. Then, the remaining chickens were sampled randomly to reach a quantity of 3. One percent of 2-mercaptoethanol (Sigma-Aldrich) in PBS was used to wash fecal contents, and samples were immediately frozen at −80°C. The intestinal tissues (duodenum to ileum) were inspected for NE lesions and scored following the criteria described by Keyburn [42], with a range of 0 (no gross lesions), 1 (congested intestinal mucosa), 2 (small focal necrosis or ulceration; one to five foci), 3 (focal necrosis or ulceration; 6 to 15 foci), and 4 (focal necrosis or ulceration; 16 or more foci). Chickens with lesion scores reaching 2 or higher were identified as NE cases, and the highest score in their small intestinal sections (duodenum, jejunum, and ileum) was recorded as the final score of NE.

### DNA extraction

Total genomic DNA was isolated from approximately 250 mg of fecal contents using the MOBIO PowerFecal^®^ DNA Isolation Kit (Mobio, Germantown, Maryland, USA) following the manufacturer’s protocol with some modifications. After adding bead solution and lysis buffer, the mixture was heated in a water bath at 65°C for 30 minutes followed by 5 minutes of vortexing. The concentration and quality of harvested DNA were determined by NanoDrop™ One Microvolume UV-Vis Spectrophotometer (Fisher Scientific) and visualized on 0.8% agarose gel (BD Biosciences, San Jose, California, USA). Afterward, genomic DNA was stored at −20°C until further analysis.

### 16S rRNA library preparation and sequencing

The variable V3-V4 region of the 16S rRNA gene was PCR-amplified in 25-μl reaction mixtures, containing 12.5 μl Clontech Labs 3P CLONEAMP HIFI PCR PREMIX (Fisher Scientific), 1 μl of each 10-μm Illumina primer (forward primer-5’CCTACGGGNGGCWGCAG 3’ and reverse primer-5’ GACTACHVGGGTATCTAATCC 3’) with standard adapter sequences, and 1 μl of DNA template. The PCR conditions started with an initial denaturation step at 95°C for 3 minutes, followed by 25 cycles of 95°C for 30 seconds, 55°C for 30 seconds, and 72°C for 30 seconds, and a final extension step at 72°C for 5 minutes on Applied Biosystems GeneAmp PCR System 9700 (Applied Biosystems Inc., Foster City, California, USA). The amplicons were cleaned up by Monarch^®^ DNA Gel Extraction Kit (New England Biolabs, Ipswich, Massachusetts, USA). Subsequently, an index PCR was performed by using Nextera XT Index Kit (Illumina, San Diego, California, USA) to attach a unique 8-bp barcode sequence to the adapters. The applied 25-μl reaction was composed of 12.5 μl KAPA HiFi HotStart Ready Mix (Kapa Biosystems, Wilmington, Massachusetts, USA), 2.5μl of each index primer, and 1 μl of 16S rRNA amplicon and reaction conditions were as follows: 95°C for 3 minutes, 8 cycles of 95°C for 30 seconds, 55°C for 30 seconds, 72°C for 30 seconds, and 72°C for 5 minutes on Mastercycler^®^ pro (Eppendorf AG, Hamburg, Germany). The PCR products were purified using Agencourt AMPure XP beads (Beckman Coulter, Indianapolis, Indiana, USA), and the size and concentration were determined by Bioanalyzer with DNA 1000 chip (Agilent, Santa Clara, California, USA) and Qubit^®^ 2.0 Fluorometer with Qubit™ dsDNA HS Assay Kit (Fisher Scientific). Those libraries were normalized and pooled to one tube with the final concentration of 10 pM. Samples were thereafter sequenced on the MiSeq^®^ System using Illumina MiSeq Reagent Kit v3 (2×300 bp paired-end run).

### Sequence processing and data analysis

Paired-end sequences were merged by means of fast length adjustment of short reads (FLASH) v1.2.11 [43] after trimming of primer and adapter sequences. Reads were de-multiplexed and filtered by Quantitative Insights into Microbial Ecology (Qiime) software v1.9.1 [44], meeting the default quality criteria and a threshold phred quality score of Q ≥ 20. Chimeric sequences were filtered out using the UCHIME algorithm [45]. The pick-up of operational taxonomic units (OTUs) was performed at 97% similarity by the UPARSE algorithm [46] in USEARCH [47].

The OTUs were further subjected to the taxonomy-based analysis by RDP Classifier v2.11 with a cut-off of 80% [48] using the Silva v128 database. Differential abundance of OTU among treatments was evaluated by metagenomeSeq. The clustered OTUs and taxa information were used for diversity and statistical analyses by Qiime v1.9.1 and R package v.3.3.1 (http://www.R-project.org/). Differences of taxonomic profiles between groups were compared using Statistical Analysis Metagenomic Profiles (STAMP) software [49] v2.1.3 with Welch’s t-test. Furthermore, LEfSe (linear discriminant analysis effect size) from the LEfSe tool (http://huttenhower.sph.harvard.edu/lefse/), an algorithm for high-dimensional class comparisons between biological conditions, was used to determine the significant feature taxa between groups or intestinal location. It emphasizes statistical significance, biological consistency, and effect relevance and allows researchers to identify differentially abundant features that are also consistent with biologically meaningful categories [50]. The Kruskal-Wallis rank sum test was included in LEfSe analysis to detect significantly different abundances and performed LDA scores to estimate the effect size (threshold: ≥ 4).

## Results

### NE reproduction and effects of lauric acid as an alternative prevention

Six of the NE cases were identified in three CP1-challenged groups (**Table 1**). They showed different degrees of characteristic gross lesions in small intestinal tissues. The most severe lesions were found in the jejunum, between its proximal end and Meckel’s diverticulum. Under co-infection with CP1 and *Eimeria*, the incidence and severity of NE increased. No NE mortality was noticed. Statistically significant differences of lesion score (LS) were determined between three CP1-challenged groups (A, B, and C) and the control counterpart (*p ≤ 0.05*). The co-infection groups (B and C) demonstrated a highly significant difference (*p ≤ 0.01*). However, the supplementation of lauric acid did not reduce the incidence and severity which were similar to the NE positive control group.

**Table 1.** NE frequency and mean lesion score by groups.

| Group | Treatment | NE lesion score |  |  |  |  | Subtotal | Lesion score | NE case |
| --- | --- | --- | --- | --- | --- | --- | --- | --- | --- |
|  |  | 0 | 1 | 2 | 3 | 4 |  |  |  |
| A | CP1 | 0 | 9 | 1 | 0 | 0 | 10 | $1.11 \pm 0.31^a$ | 1 |
| B | CP1+*Eimeria* | 0 | 8 | 0 | 1 | 1 | 10 | $1.50 \pm 1.02^a$ | 2 |
| C | CP1+*Eimeria*+LA | 0 | 7 | 1 | 1 | 1 | 10 | $1.60 \pm 1.02^a$ | 3 |
| D | *Eimeria* | - | - | - | - | - | 10 | - | 0 |
| E | CTL | 5 | 4 | 0 | 0 | 0 | 9 | $0.44 \pm 0.50^b$ | 0 |

LA lauric acid; NE necrotic enteritis; CTL: control group.
NE case: lesion score reaching 2 or above.
One chick in CTL was misclassified during the trial and excluded.
Dissimilar letters indicate a significant difference at a level of $\alpha=0.05$ .

### Metadata and sequencing

A total of 11,191,102 sequence reads with an average length of 453 ±5 base pairs were obtained from 30 samples, including 15 jejunal samples (3 samples per group in AJ, BJ, CJ, DJ, and EJ) and 15 cecal samples (3 samples per group in AC, BC, CC, DC, and EC). The sequences were filtered and further clustered into OTU using a cut-off of 97% similarity. The estimate of Good’s coverage reached 98% for all the jejunal and cecal samples. The rarefaction curve demonstrated that the sequencing depth was adequate to cover the bacterial diversity in the jejunal and cecal samples (**Figure S1**).

### Normal microbial composition in the jejunum and cecum

*Firmicutes* (92.1% of relative abundance) was the most dominant phylum in the jejunum, followed by *Cyanobacteria* (2.2%) and *Proteobacteria* (2.1%), *Bacteroidetes* (1.9%), and *Actinobacteria* (1.7%). On the contrary, the phylum of *Bacteroidetes* (75.5%) predominated in the cecum, followed by *Firmicutes* (19.8%) and *Proteobacteria* (4.7%) (**Figure 1A**). At the genus level, jejunal contents were dominated by *Lactobacillus* (41.2% of relative abundance) and *Clostridium sensu stricto 1* (39.1%), followed by other unclassified genus (8.7%), *Weissella* (3.6%), *Enterococcus* (1.9%), *Escherichia Shigella* (1.8%), and *Staphylococcus* (1.6%). *Bacteroides* (75.5%) was the most abundant genus in the cecum, followed by other unclassified genus (17.2%), *Escherichia Shigella* (3.1%), *Eisenbergiella* (1.7%), and *Anaerotruncus* (1.5%) (**Figure 1B**). The genera of *Lactobacillus*, *Clostridium sensu stricto 1*, *Weissella*, *Enterococcus*, *Staphylococcus*, and *Bifidobacterium* in the jejunum exhibited significant difference in abundance compared to those in the cecum. Cecal microbiota contained significantly higher abundance of *Bacteroides* and *Proteus* (Welch’s t test, *p < 0.05;* **Figure S2**).

**Figure 1.**
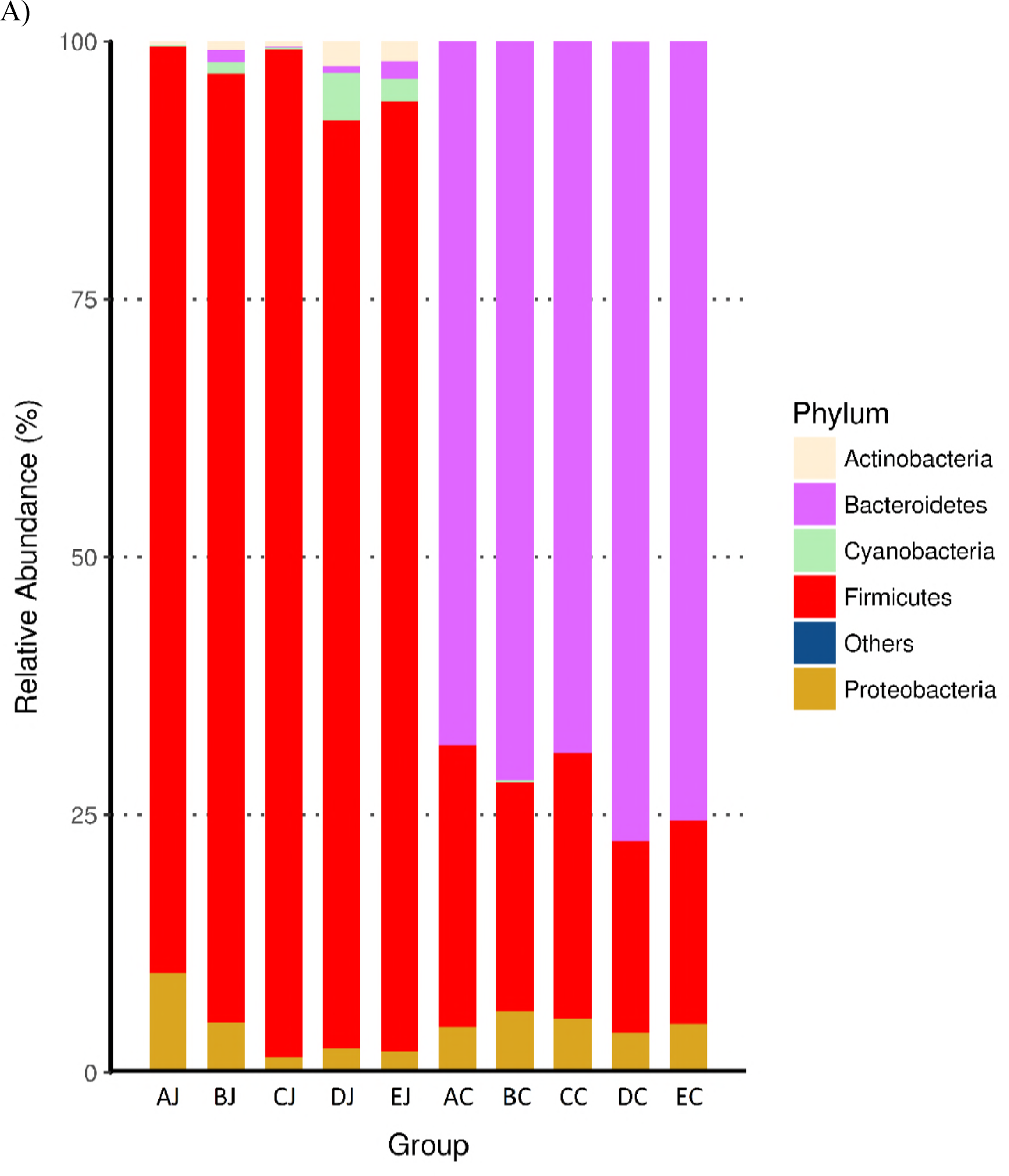
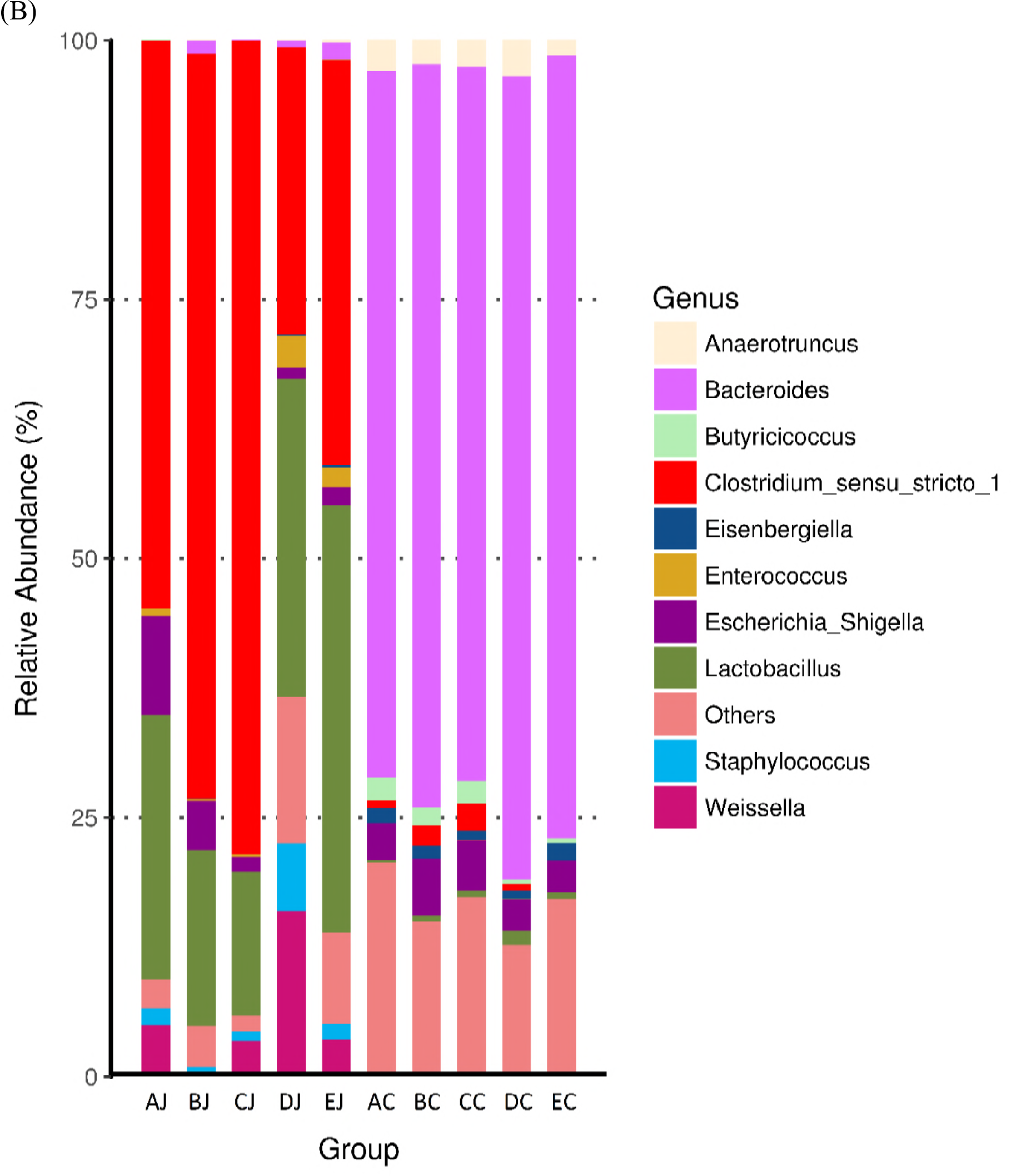
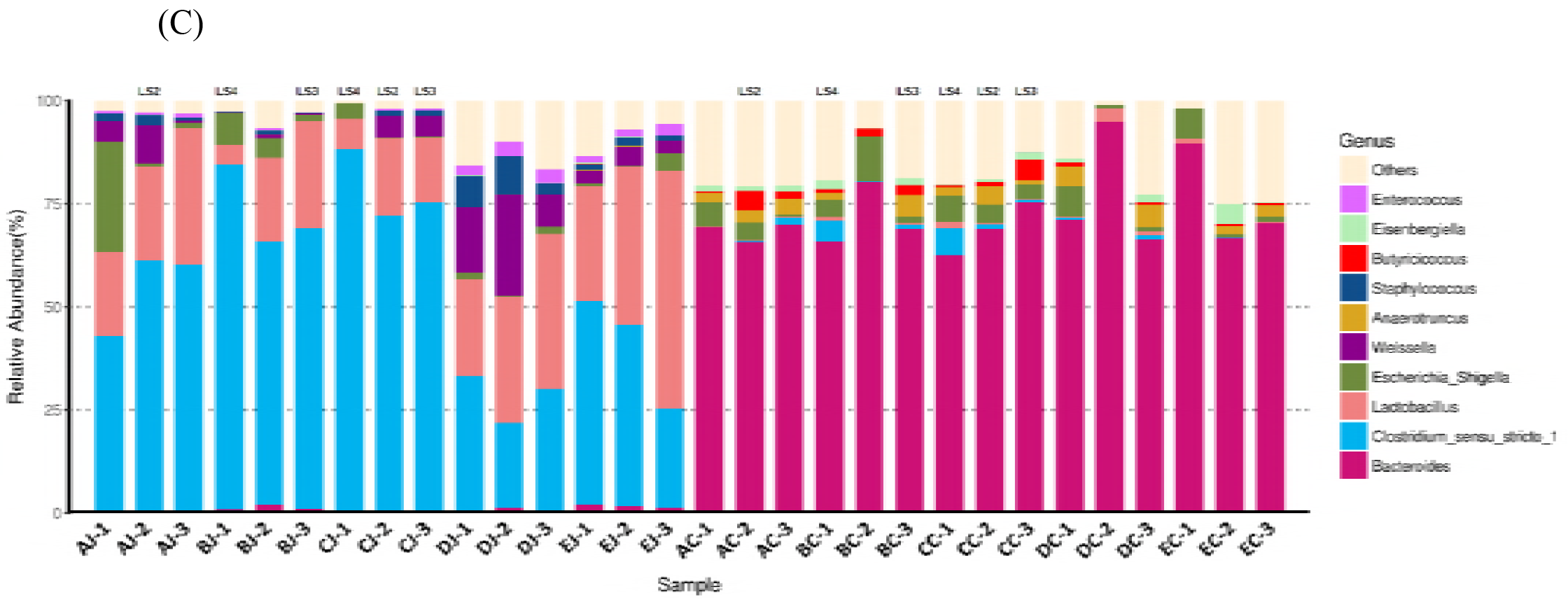
Microbiota composition in jejunum and cecum with different treatments. Each bar represents the average relative abundance of each bacterial taxon within a group. The top 5 and 10 abundant taxa are shown at the level of phylum and genus, respectively. (A) Abundant phyla in jejunum and ceca by groups; (B) Abundant genera in jejunum and ceca by groups; (C) Abundant genera in jejunal and cecal samples. LS stands for lesion score of NE.

### Changes of microbial communities in response to treatments

In the jejunum, challenge of CP1 increased the relative abundance of the genera of *Clostridium sensu stricto 1* (54.75%), *Escherichia Shigella* (9.57%), and *Weissella* (4.99%) but significantly decreased the population of *Lactobacillus* (25.44%) (**Figure 2**). The inoculation of *Eimeria* to chickens significantly increased the relative abundance of *Weissella* (16.01%) and *Staphylococcus* (6.51%), but decreased the amount of *Lactobacillus* (30.66%) and *Clostridium sensu stricto 1* (27.69%). Co-infection with CP1 and *Eimeria* led to significant increment of *Clostridium sensu stricto 1* (71.89%), increased relative abundance of *Escherichia Shigella* (4.68%), but the decrements of *Lactobacillus* (16.99%)*, Weissella* (0.44%) and *Staphylococcus* (0.40%). In the cecum, different treatments did not promote significant difference of taxa abundance between groups with an exception of *Eisenbergiella, significantly increased in* co-infection group. However, challenge of CP1 and co-infection of CP1 and *Eimeria* still promoted cecal increments of *Clostridium sensu stricto 1* (the relative abundance of this taxon in groups challenging with CP1, CP1 and *Eimeria*, and control was 0.75%, 1.99%, and 0.02%).

**Figure 2.**
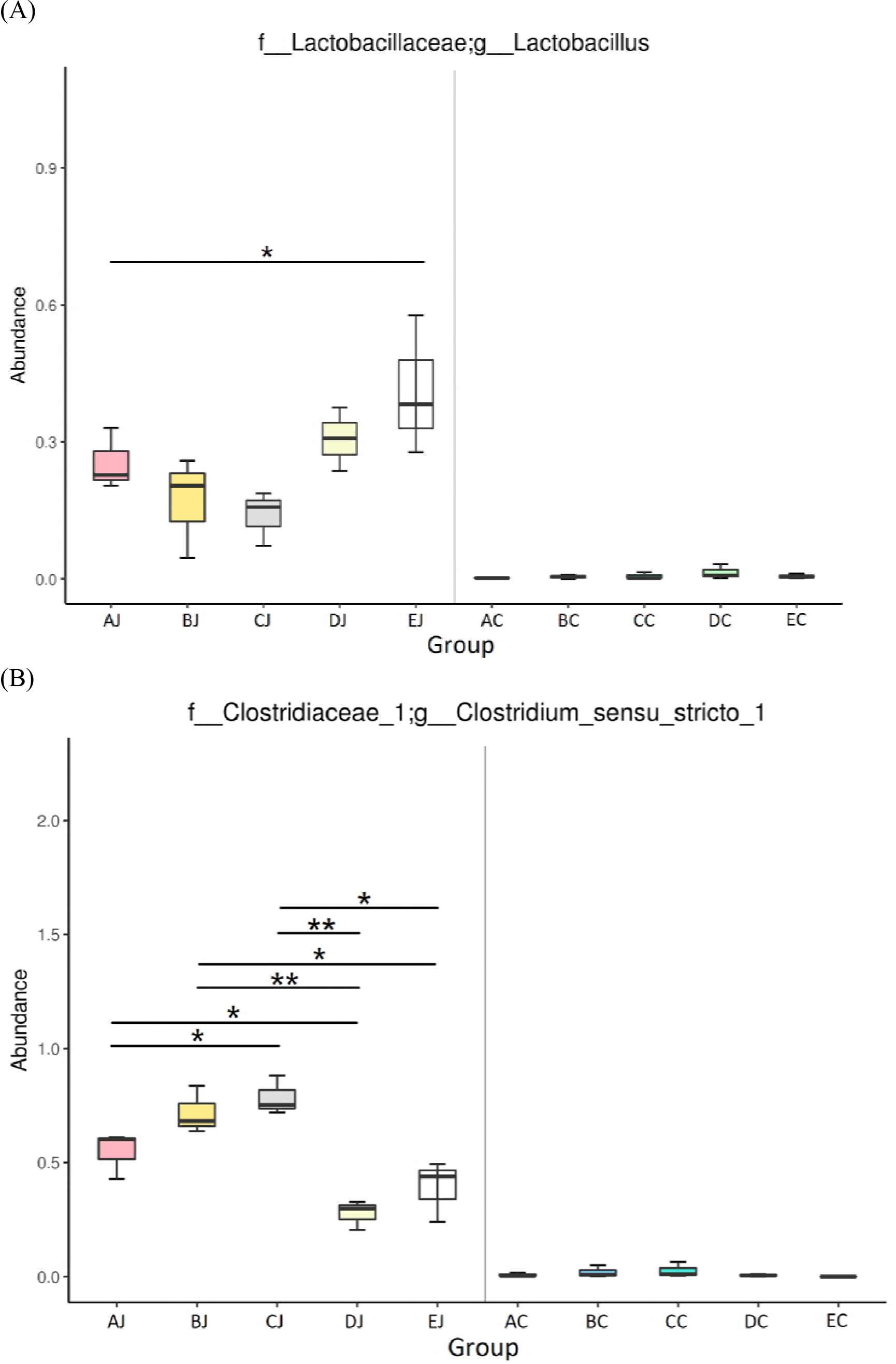
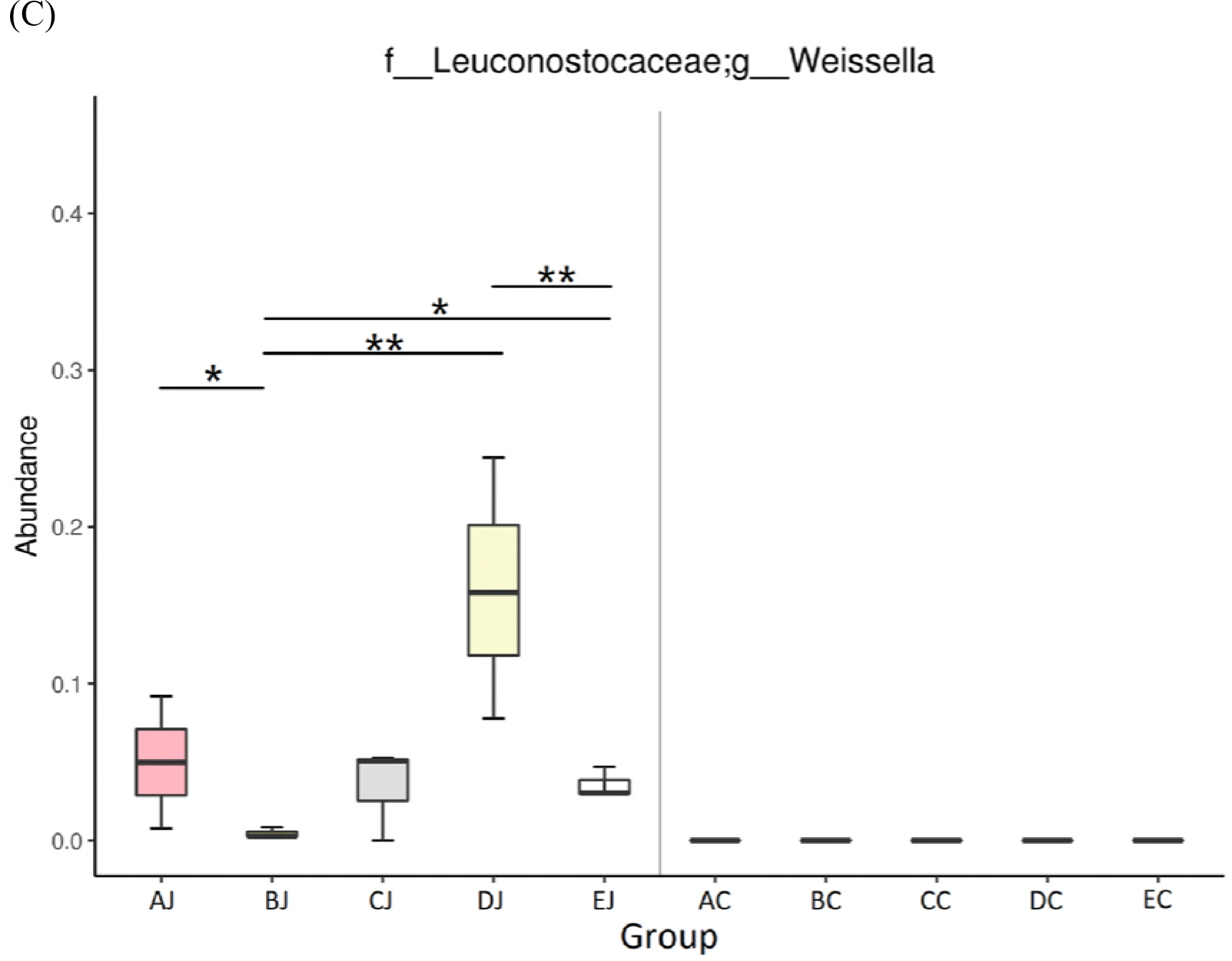
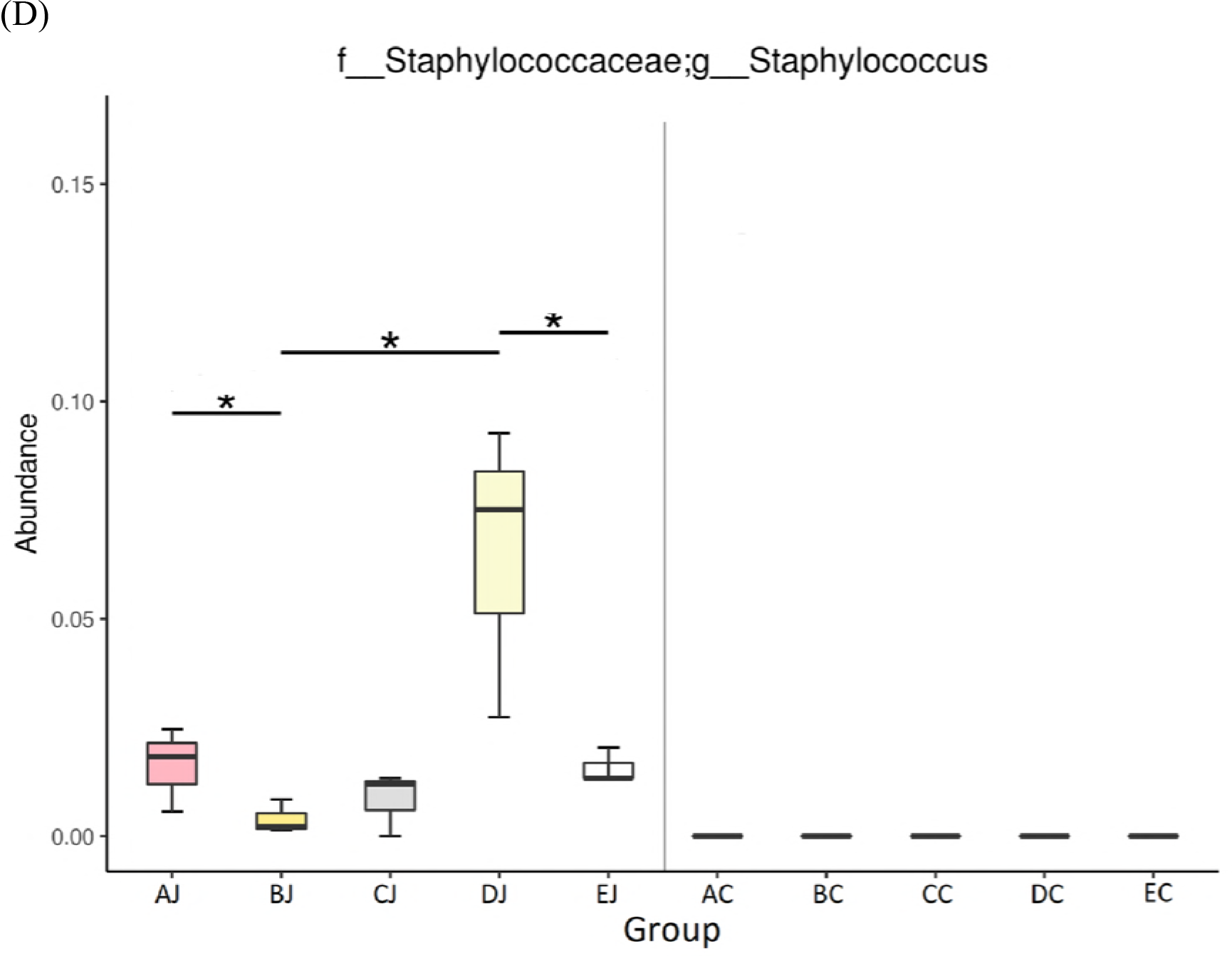
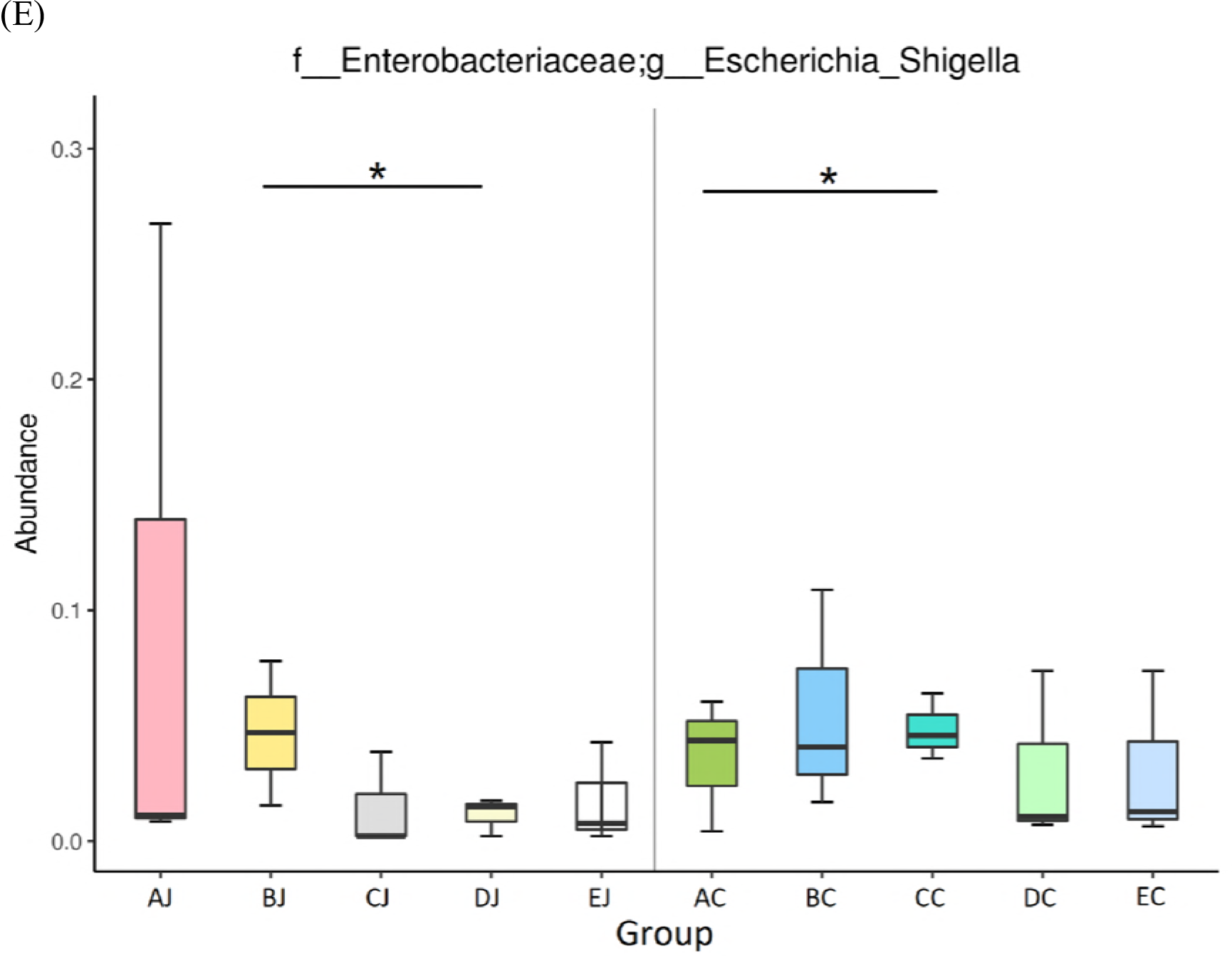
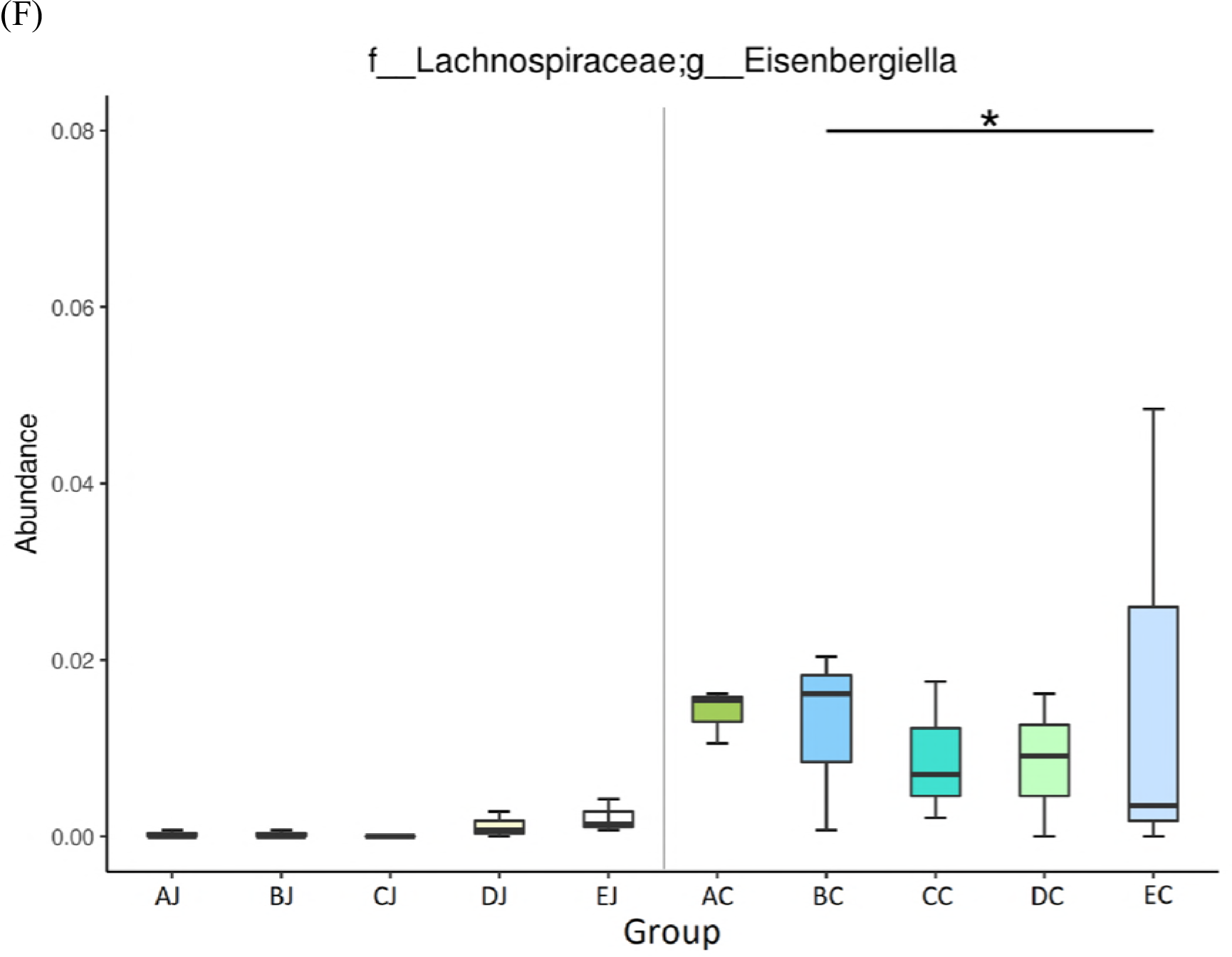
Differential abundance of genera between groups by metagenomeSeq. (A) *Lactobacillus*; (B) *Clostridium sensu stricto 1*; (C) *Weissella*; (D) *Staphylococcus*; (E) *Escherichia Shigella*; (F) *Eisenbergiella*. * *p ≤ 0.05* and ** *p ≤ 0.01*.

### Microbial diversities in response to treatments

In jejunal microbiota, challenge of CP1 (AJ) and co-infection with CP1 and *Eimeria* (BJ) reduced species richness and evenness, but the infection of *Eimeria* (DJ) exerted counter results. Addition of lauric acid into co-infection group (CJ) exacerbated the reduction observed in BJ group. However, no apparent effect was noted on cecal microbiota following above treatments (**Figure S3 and Figure 3**). Analysis of alpha diversity by Shannon index further demonstrated that challenge of CP1 in conjunction with *Eimeria* infection significantly reduced species diversity in jejunal microbiota. The 16S rRNA gene survey by principal coordinate analysis (PCoA) and principal component analysis (PCA) showed a distinct separation of two community profiles between the jejunal and cecal microbiota. Cluster and heat map analyses exhibited distinct classifications and microbial compositions between the jejunum and cecum, coinciding with observations on PCoA and PCA (**Figure 4)**. Additionally, the results of PCoA and PCA also depicted the differential diversity between the CP1-challenged (group AJ, BJ and CJ), *Eimeria*-infected (DJ), and control (EJ) groups in jejunal microbiota, showing that challenge of CP1 shared similar microbial community structures with co-infection with CP1 and *Eimeria*. However, cecal groups with CP1 treatments did not display cluster phenomenon as jejunal groups displayed in PCoA. PCA with hierarchical clustering further reflected that *Clostridium sensu stricto 1* was contributory to the similarity of NE assemblage, and the genera of *Lactobacillus, Weissella*, and *Staphylococcus* contributed to discrepant community structures in *Eimeria*-treated and control groups in the jejunum. On the other hand, *Bacteroidetes* was the main genus contributing to the distinct separation between jejunal and cecal groups (**Figure 5**).

**Figure 3.**
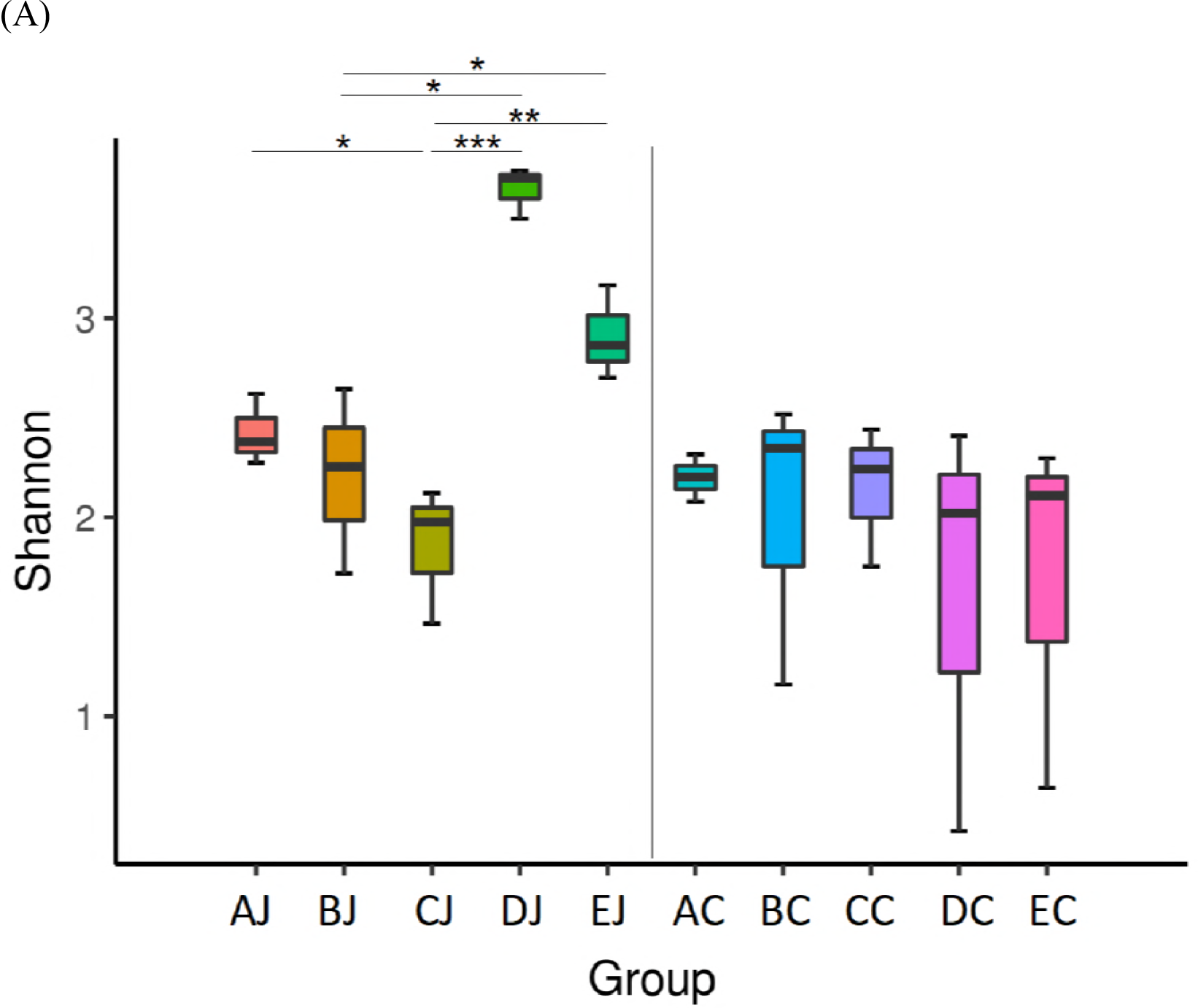
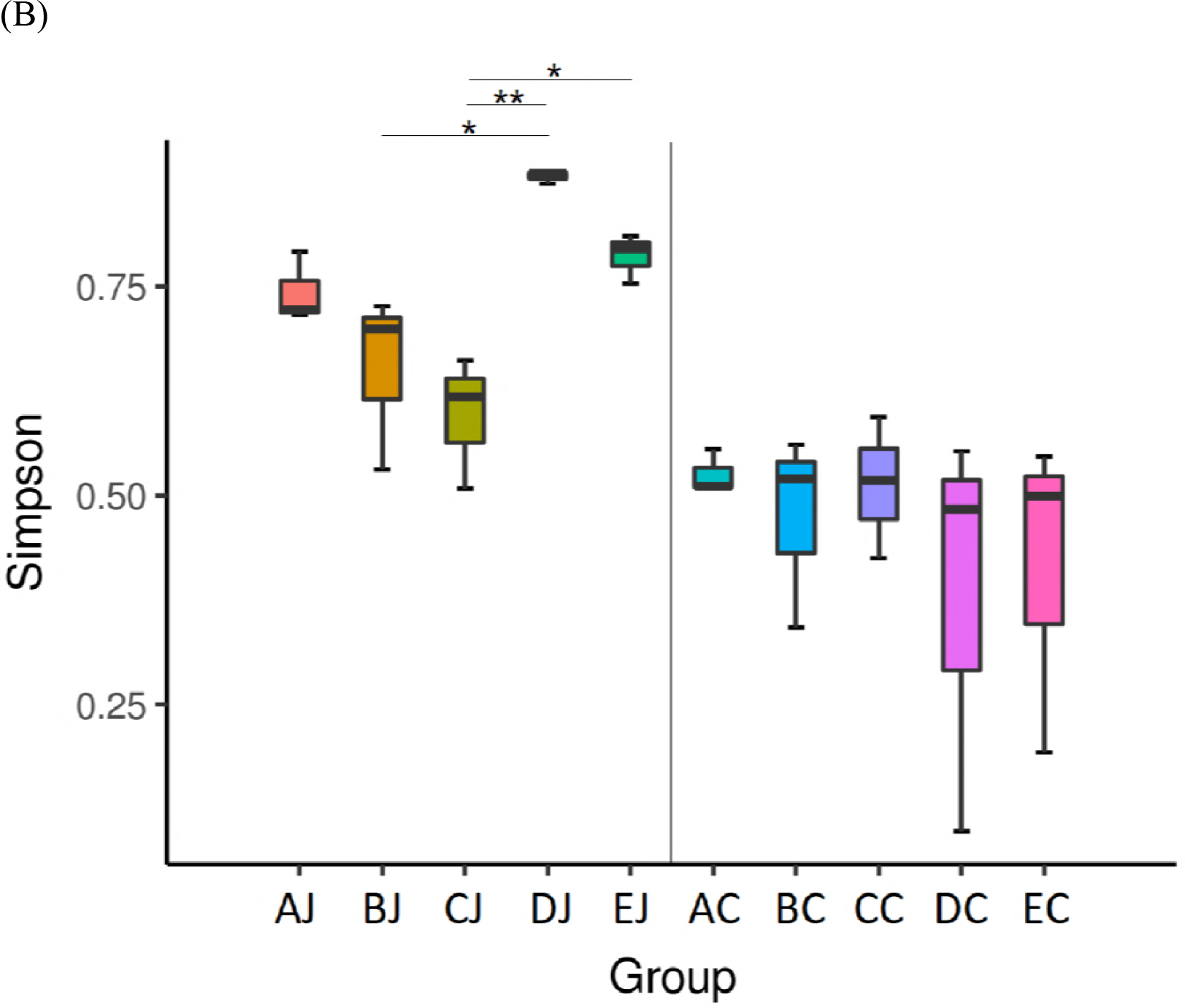
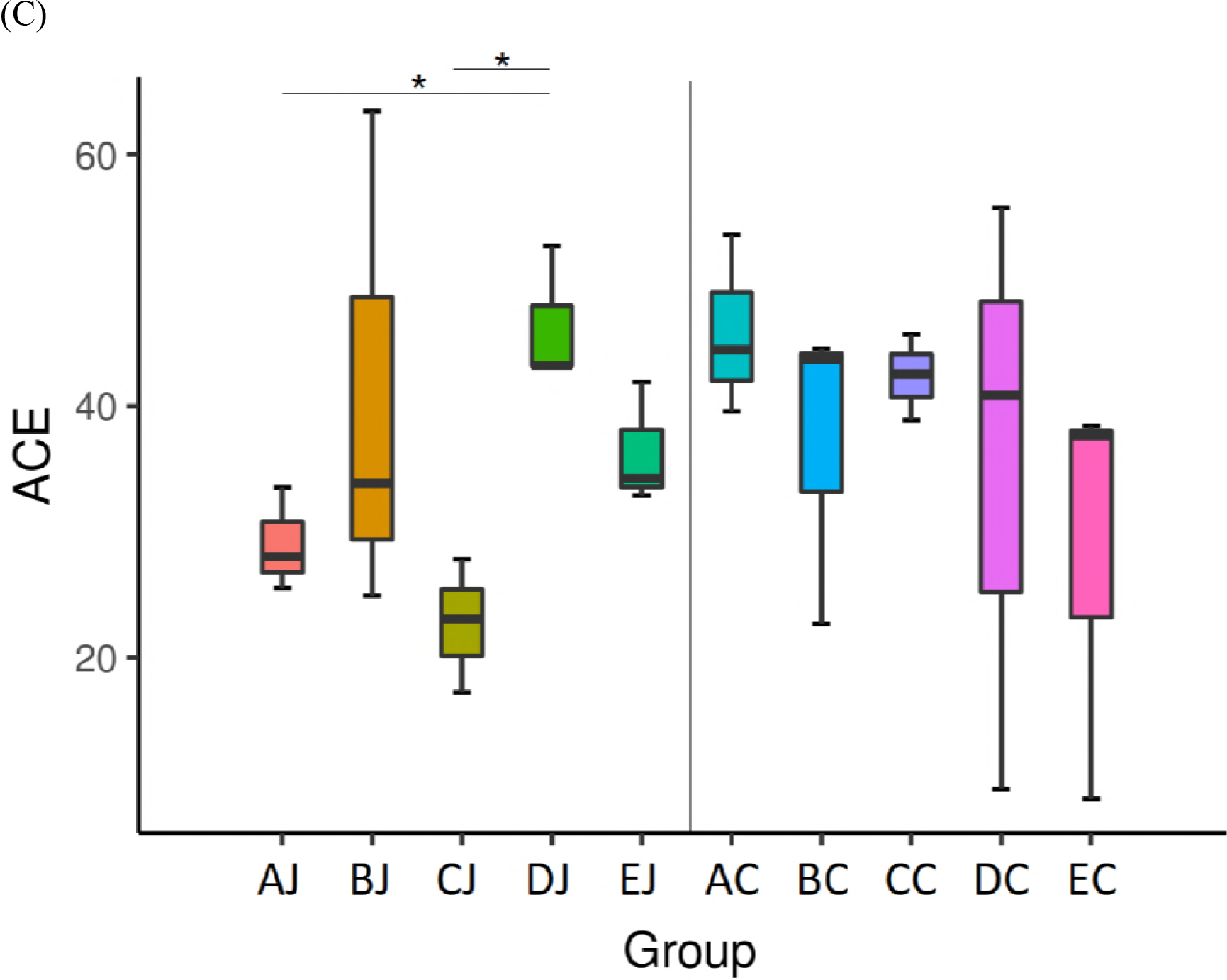
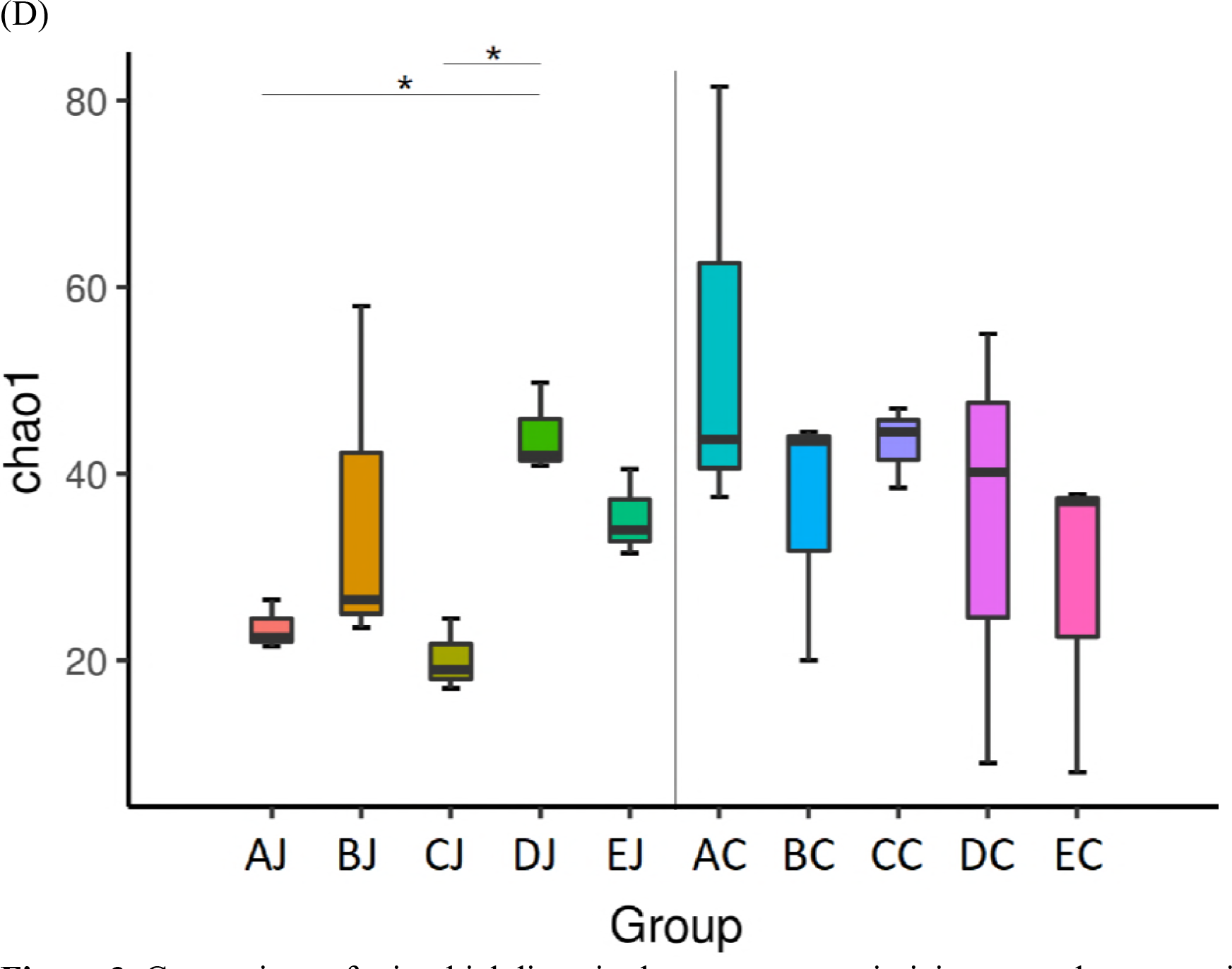
Comparison of microbial diversity between groups in jejunum and cecum using different measures of alpha diversity. (A) Shannon index, (B) Simpson index (C), abundance-based coverage estimator (ACE) index, and (D) Chao1 index. Results are shown as mean ± SEM. Kruskal-Wallis test: * *p ≤ 0.05*, ** *p ≤ 0.01, and *** p ≤ 0.001*.

**Figure 4.**
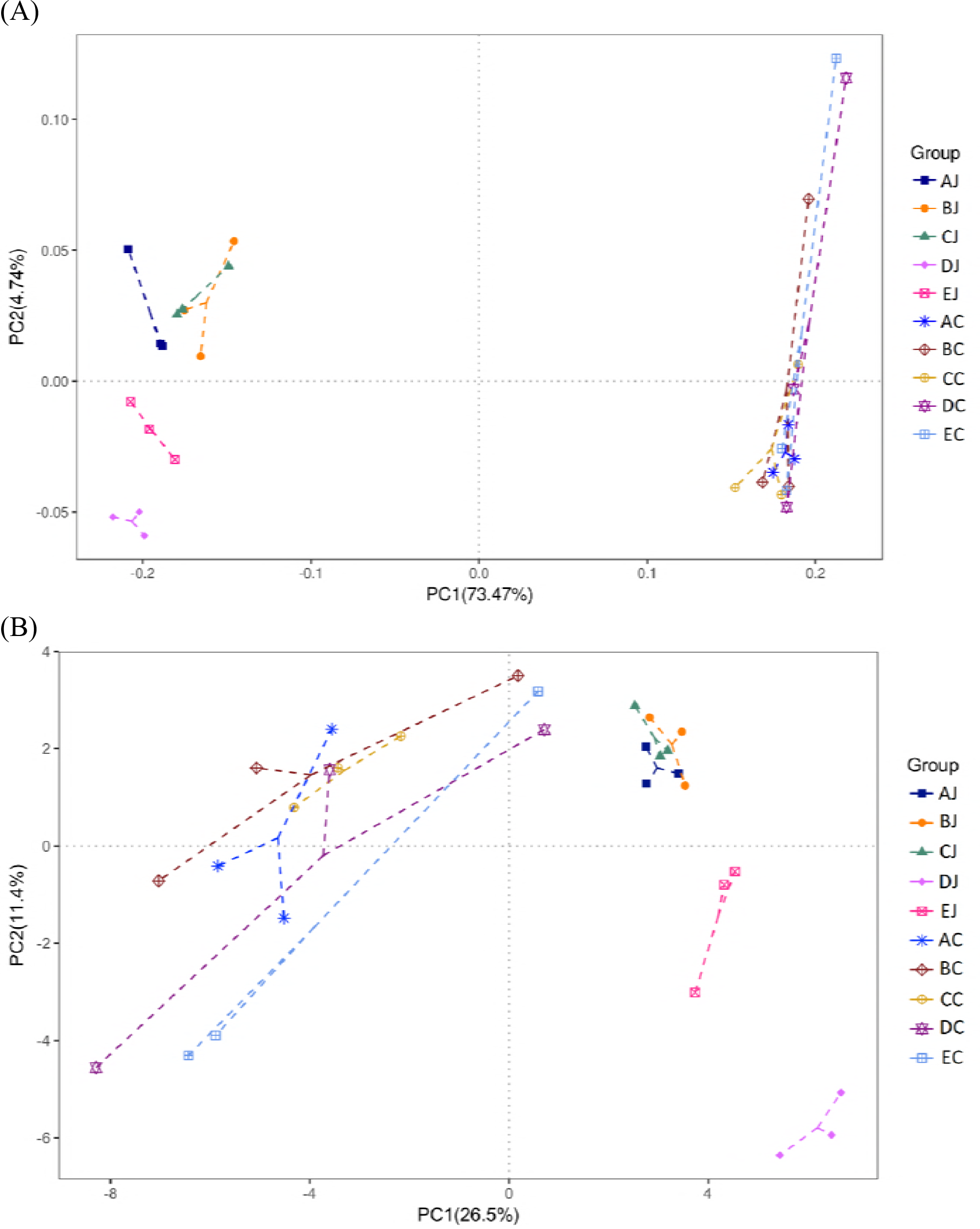
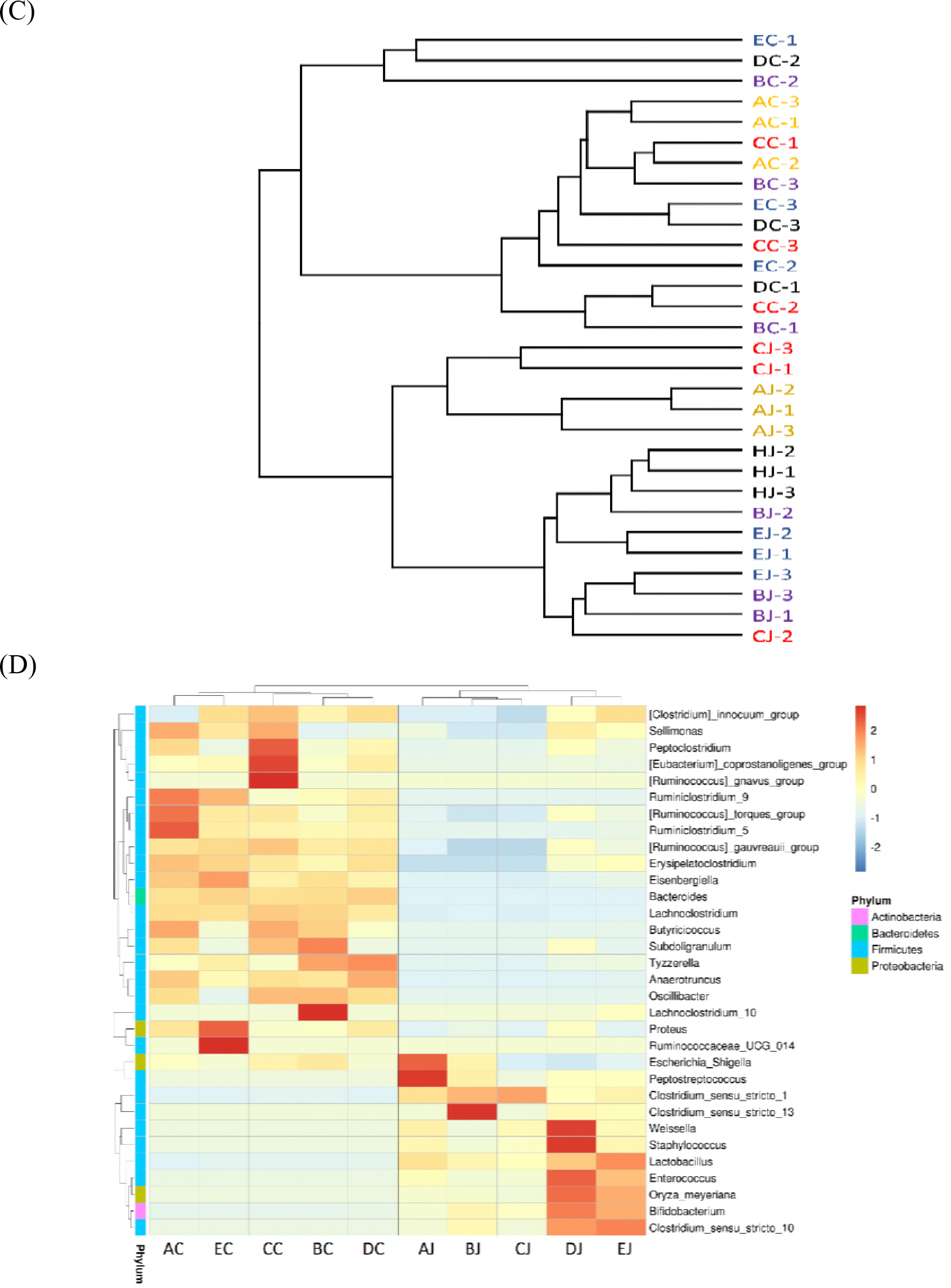
Comparison of the compositions and similarities of jejunal and cecal microbiota with different treatments. (A) Weighted Unifrac principal coordinate analysis (PCoA); (B) principal component analysis (PCA); (C) cluster analysis by unweighted paired-group method using arithmetic means (UPGMA) using unweighted Unifrac distance; (D) heat map analysis at the genus level.

**Figure 5.**
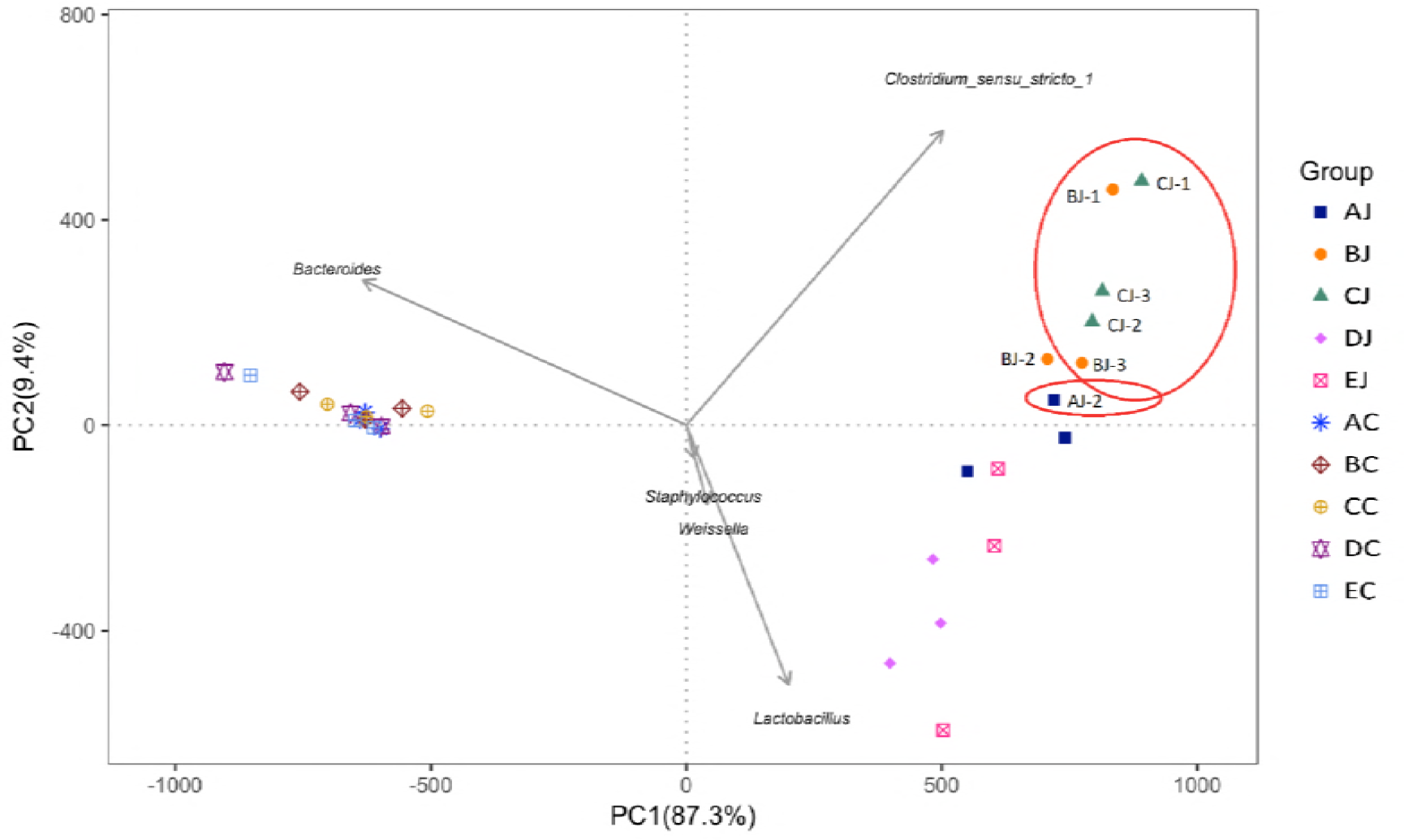
Principal component analysis and hierarchical clustering of contributory genus to NE assemblage (in red circle) and to the dissimilarity between groups.

### Microbial community structure and taxa contributory to NE

Analysis of jejunal microbiota in NE cases revealed that *Clostridium sensu stricto 1*, to which causative *C. perfringens* belongs, was the most dominant genus, followed by *Lactobacillus, Weissella*, *Escherichia Shigella*, *Staphylococcus*, and others. Accompanying the elevation of NE severity, the relative abundance of *Clostridium sensu stricto 1* increased (relative abundance ≥ *75%* in LS4 compared to 50-75% in LS2 and LS3). Conversely, the population of *Lactobacillus* decreased while the lesion score was elevated. The relative amount of *Escherichia Shigella* was variable in NE cases, presenting higher abundance after CP1 challenge but low population following co-infection with CP1 and *Eimeria*. (**Figure 1C**).

Heat map analysis exhibited that NE cases harbored the similar microbial community profile. *Clostridium sensu stricto 1* and *C. perfringens* were consistently presented and abundant taxa in jejunum (**Figure 6**). Opposite low abundance of *Lactobacillus* was noted. However, only the increment of *Clostridium sensu stricto 1* but not *C. perfringens* (data not shown) demonstrated significance on NE by metagenomeSeq (**Figure 2B**). Using Welch’s t-test, jejunal groups further showed that CP1 in conjunction with *Eimeria* increased significantly *Clostridium sensu stricto 1* and *C. perfringens* when compared to the control (**Figure S4**; *p < 0.05)*, whereas challenge of CP1 alone did not lead to significant increase of these taxa. Differential abundant phylotypes between different treatments in jejunum were further evaluated by LEfSe using the LDA score of 4. This threshold guarantees that the meaningful taxa is compared and eliminates most of rare taxa. LEfSe demonstrated similar results as Welch’s test that challenge of CP1 unable to yield a significantly higher amount of *Clostridium sensu stricto 1* and *C. perfringens*. However, significant differences were displayed when CP1 co-infected with *Eimeria* (**Figure 7**). No differential taxon was found in cecal groups (AC, BC, CC, HDC, and EC) while Welch’s t-test and LEfSe were applied.

**Figure 6.**
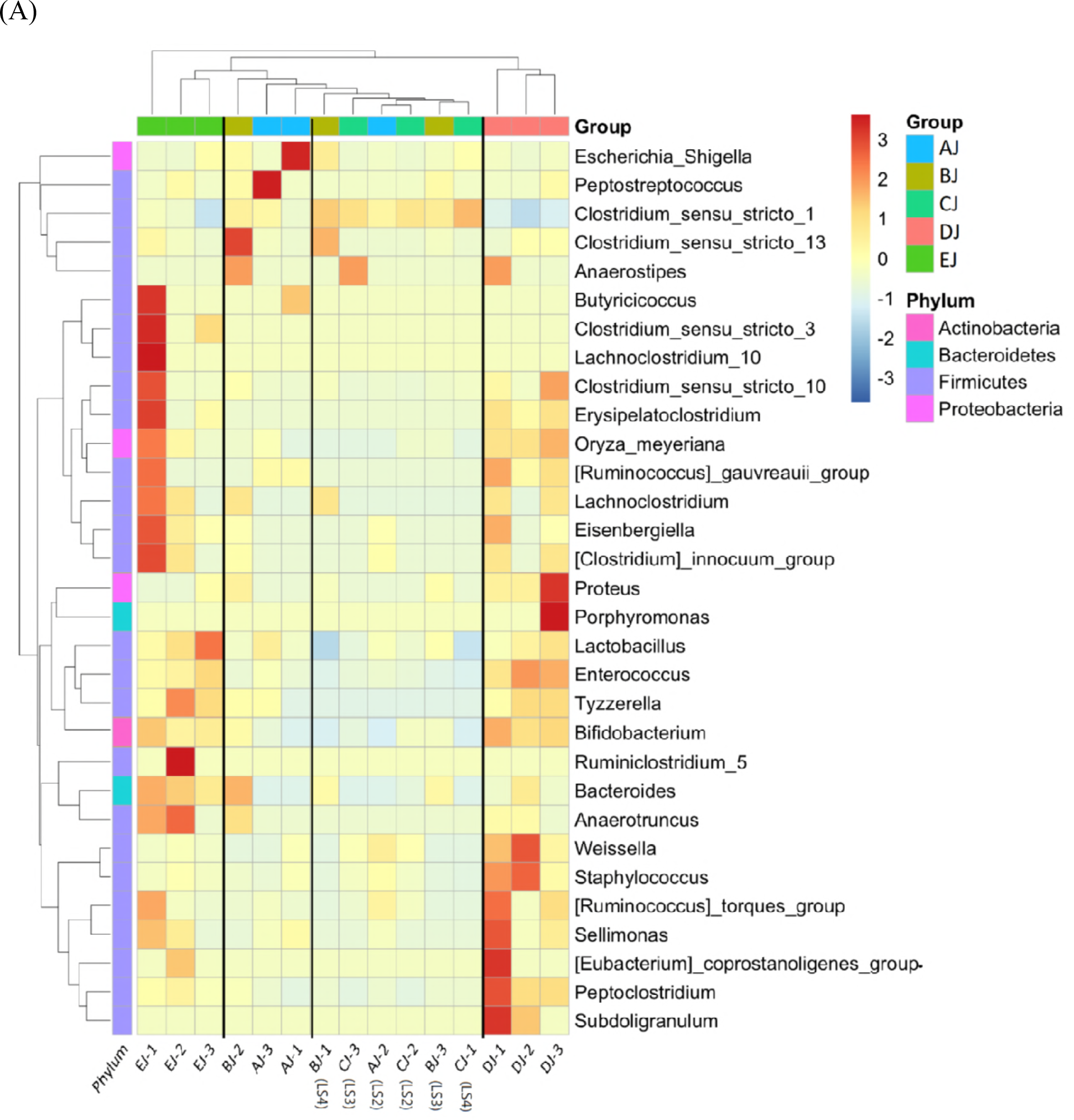
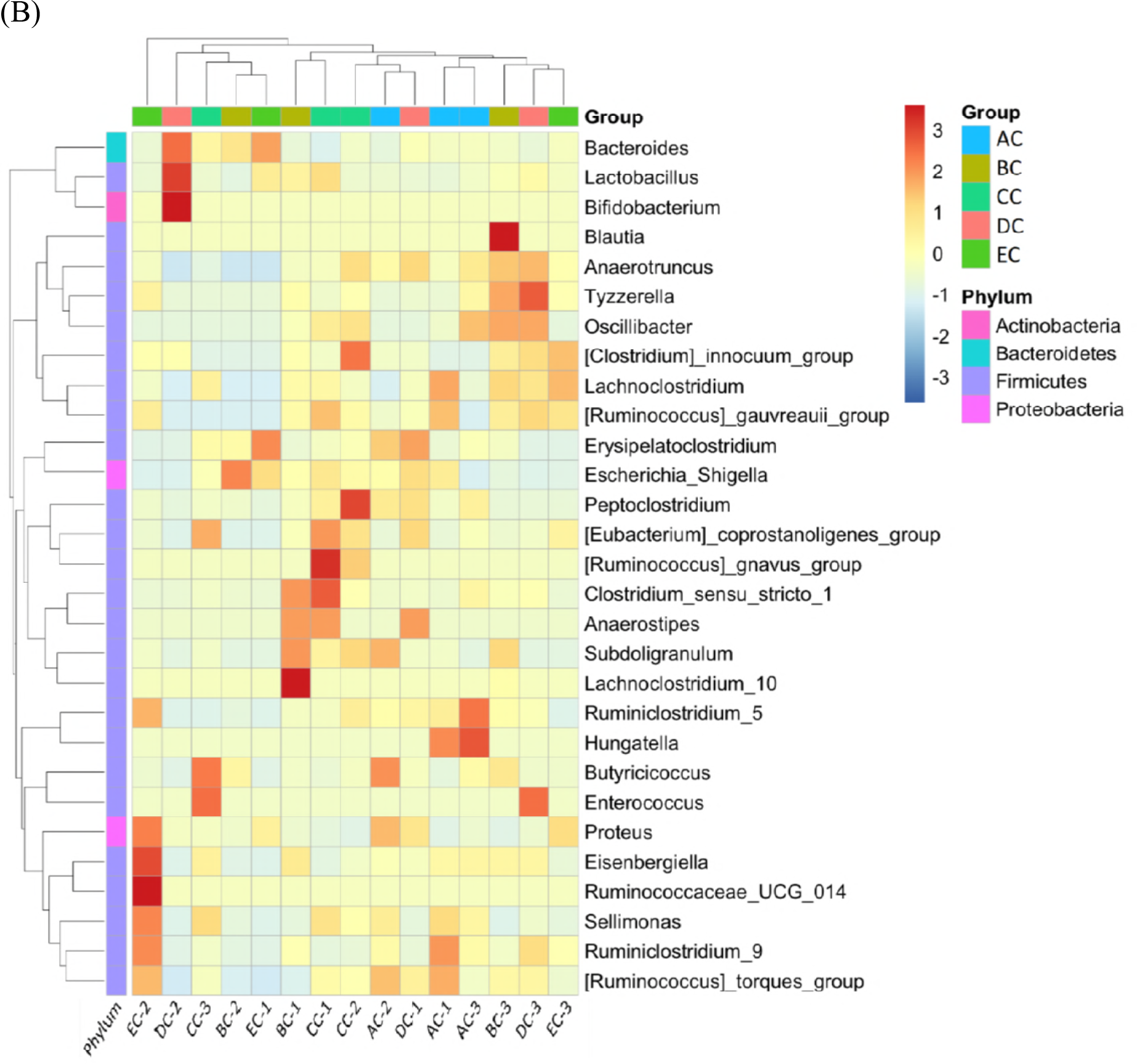
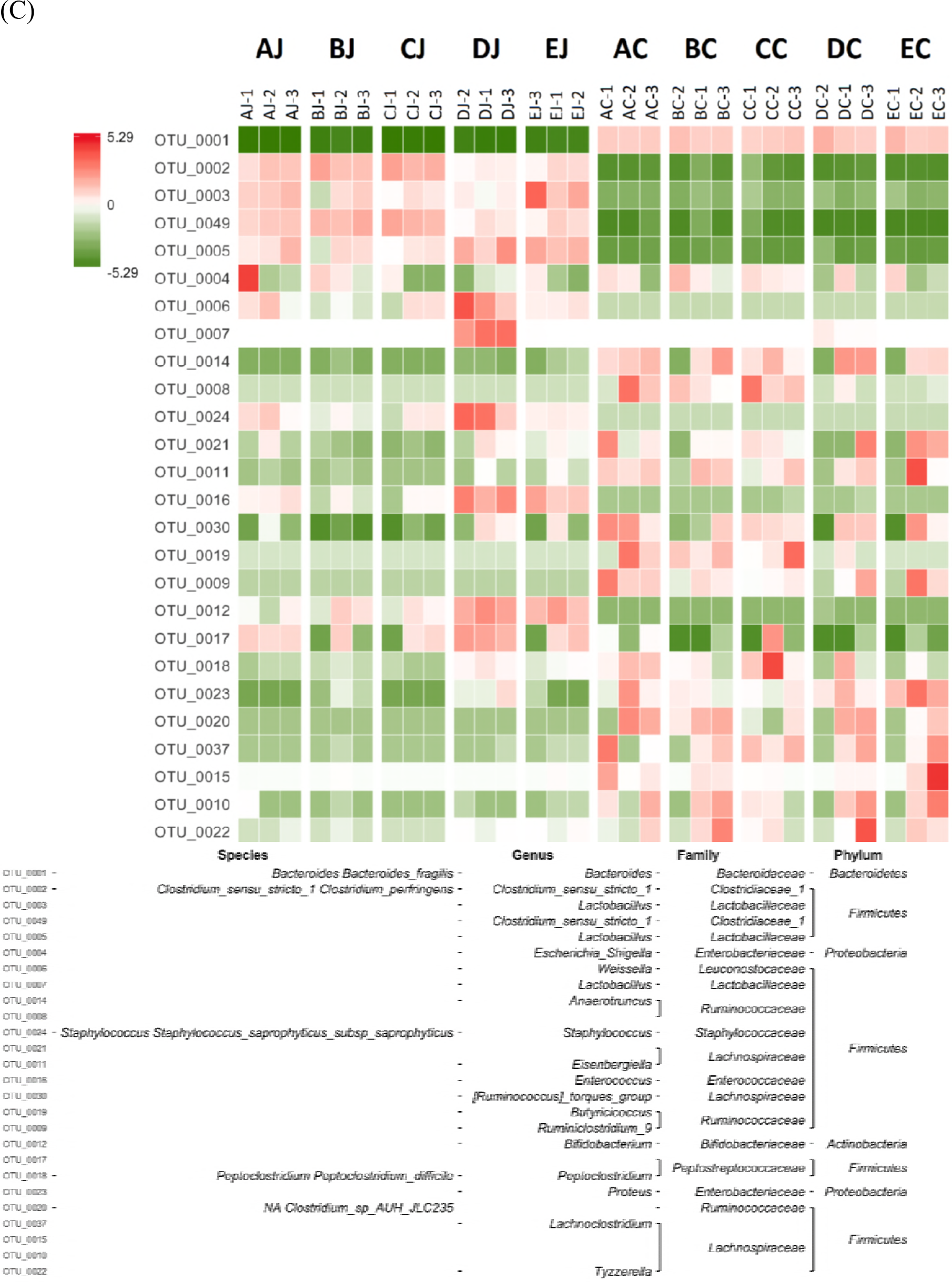
Heat map analysis of contributory taxa to NE at the genus level in jejunal (A) and cecal (B) samples. (C) Heat map of gut bacteria with the relative abundance of OTUS by z score and represented bacterial taxa information, including phylum, family, genus, and species. Top 26 taxa was shown.

**Figure 7.**
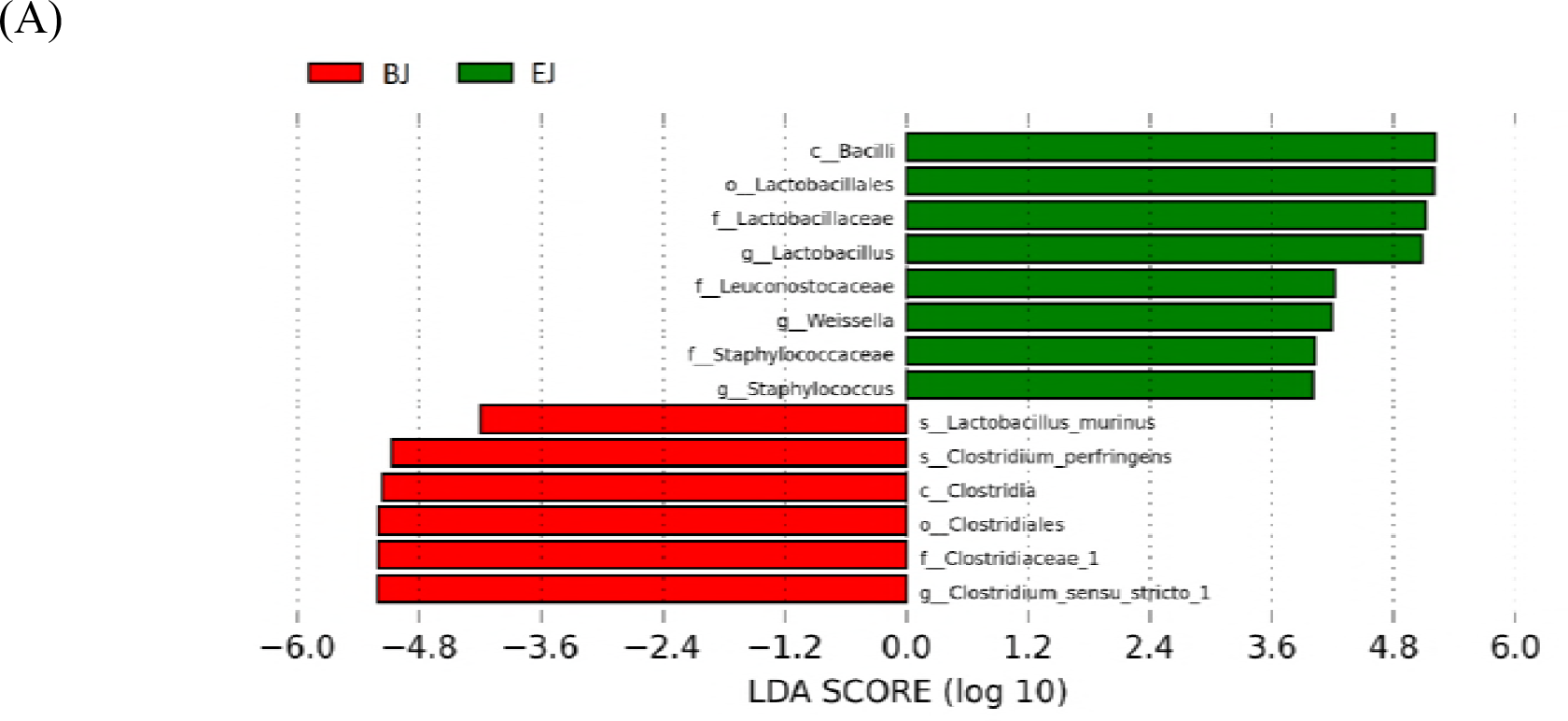
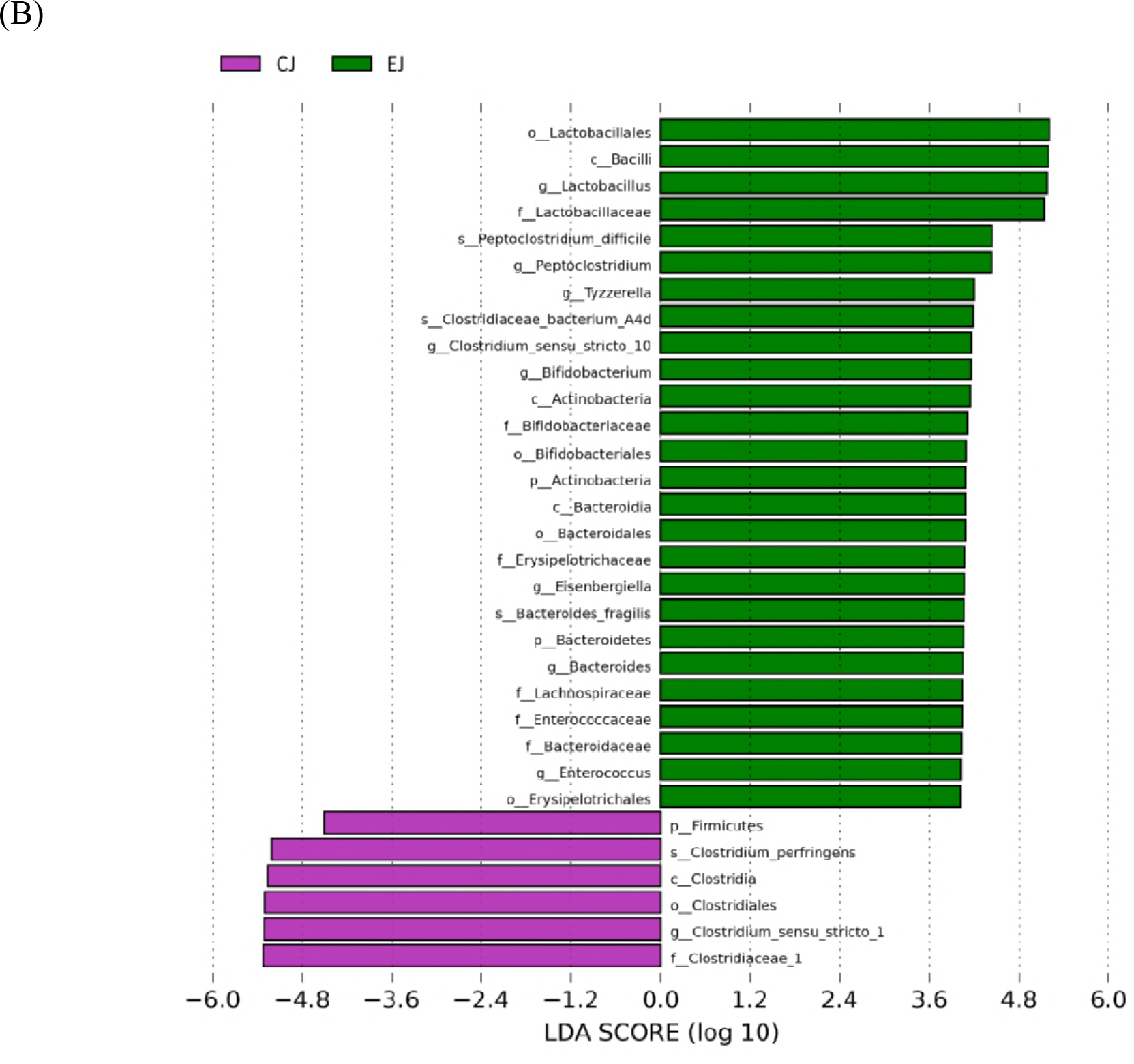
LEfSe identified the most differentially abundant clades at all taxonomic levels between jejunal groups using the LDA score of 4. Differentially abundant taxa in group BJ versus EJ (A) and CJ versus EJ (B).

### Comparison of gut metagenomes in co-infected chickens with and without lauric acid

Addition of lauric acid increased the relative abundance of *Clostridium sensu stricto 1* and *Weissella* but decreased the relative amount of *Escherichia Shigella* in the jejunum compared to the co-infection group without supplementing lauric acid. Nonetheless, no significance was detected in this comparison. In addition, supplementation of lauric acid did not apparently affect the cecal microbiota between these two groups.

## Discussion

By exploring microbial composition in normal chickens, the major microbial genera in the jejunum were *Lactobacillus* and *Clostridium sensu stricto 1*, followed by other unclassified bacteria, *Weissella*, *Enterococcus*, *Escherichia Shigella*, and *Staphylococcus*. *Bacteroides* was the most abundant group in the cecum, and the remaining taxa were sequentially other unclassified bacteria, *Escherichia Shigella*, *Eisenbergiella*, and *Anaerotruncus*. Side by side treatments of *C. perfringens* and *Eimeria* altered microbial community compositions, significantly in jejunal microbiota. In this study, challenge of CP1 increased the abundance of *Clostridium sensu stricto 1*, *Escherichia Shigella*, and *Weissella* in the jejunum, but significantly decreased the population of *Lactobacillus*. Infection of *Eimeria* significantly increased the abundance of *Weissella* and *Staphylococcus*, but decreased the amount of *Lactobacillus* and *Clostridium sensu stricto 1*. Co-infection with *C. perfringens* and *Eimeria* led to significant increment of *Clostridium sensu stricto 1*, increased abundance of *Escherichia Shigella*, but decrements of *Lactobacillus, Weissella* and *Staphylococcus.* Specifically, it decreases the α-diversity index of the small intestinal microbial community, promoting single dominance of *Clostridium sensu stricto 1* reaching the relative abundance to 71.89%. On the other hand, six NE cases shared similar microbial community profile observed in PCA, indicating there exists a certain microbiota contributory to the disease. With more NE severity, higher relative abundance of *Clostridium sensu stricto 1* but lower relative amount of *Lactobacillus* in jejunal microbiota was noted.

Several studies has been shown *C. perfringens* challenge decreased the population of *Lactobacillus* in ileum [20, 51]. Lactobacilli are known as lactic acid producing bacteria and shown to have protection at intestinal barrier by competition with pathogens. They are also able to induce immunomodulation and ferment carbohydrates into lactic acids that lower the pH of the intestinal environment to inhibit growth of acid-sensitive pathogenic bacteria [52, 53]. Therefore, suppression of lactobacilli is regularly considered beneficial to growth and colonization of enteric pathogen. This study first demonstrated the decrement of lactobacilli in jejunum following challenge of *C. perfringens* alone and in conjunction with *Eimeria*. The change of this taxon following the NE severity indicates that decrement of *Lactobacillus* may play a role in the development of NE. In addition, the increased abundance of *Escherichia Shigella* was also observed after the challenge of *C. perfringens* and co-infection with *C. perfringens* and *Eimeria*. This genus includes enteric pathogens, which can colonize in the intestines of both humans and chickens, consequently triggering specific diseases [54]. Some studies indicated that the increment of *Escherichia Shigella* in ileum was correlate with NE [55, 56]. Nevertheless, our study found that *C. perfringens* challenge could increase the abundance of *Escherichia Shigella* but the increment was not in accordance with NE occurrence. Furthermore, the reduction of this taxa abundance was noticed in lauric acid supplementing group which has higher number of NE cases. Those finding reflected a contradiction for this genus participating in NE development. Last but not least, a reduced abundance of *Weissella* in the jejunum of NE afflicted chickens was also noted. Another study reported similar result in cecal micorbiota after *C. perfringens* challenge [18]. *Weissella* are lactic acid bacteria and belong to the family of the *Leuconostocaceae*. They harbor probiotic properties and can generate several products with prebiotic potential [57]. It may interact with *C. perfringens* as other lactic acid bacteria, but its role in NE development is unclear. More studies will be needed to elucidate the relationship between *Weissella* and NE.

In current study, significant overgrowth of *Clostridium sensu stricto 1* was associated with NE and the infection of *Eimeria* precedent to challenge of *C. perfringens* exerted synergistic effects on the overrepresentation. This correlation was consistently demonstrated by analyses of metagenomeSeq, STAMP, and LEfSe. The STAMP and LEfSe further showed *C. perfringens* was significantly overrepresented in NE groups. However, such significance was not identified by metagenomeSeq when *C. perfringens* was targeted. This result indicates that, in addition to *C. perfringens*, other bacteria under the same genus of *Clostridium sensu stricto 1* also played a role in contributing to the development of disease. The *Clostridium* genus is well-classified into 19 clusters by phylogenetic analysis [58]. *Clostridium sensu stricto* are grouped around the type species *Clostridium butyricum* and belong to the *Clostridium* cluster 1within the *Clostridiaceae* family [59]. *Clostridium sensu stricto 1* contains *C. perfringens* and other real *Clostridium* species. Their members are generally perceived as pathogenic [60] as well as interpreted as an indicator of a less healthy microbiota [61]. This suggestion coincides with our finding that *C. perfringens* challenge on its own is not capable of causing significant abundance of *Clostridium sensu stricto 1* and unable to produce more NE case observed. Future research is recommended to clarify the role of other members of *Clostridium sensu stricto 1* in the pathogenesis of NE.

Single infection of *Eimeria* could not produce NE in the present study. The treatment reduced the relative abundance of *Clostridium sensu stricto 1* and *Lactobacillus*, but significantly increased *Weissella* and *Staphylococcus* in jejunal microbiota*. Eimeria* infection has been shown to provide nutrients for *C. perfringens* to grow and cause physical damage to gut epithelium, thus facilitating the colonization and proliferation of *C. perfringens* [8, 62, 63]. However, the inoculation of *Eimeria* into normal chickens did not elicit overgrowth of *Clostridium sensu stricto 1* and *C. perfringens* except challenging with exogenous *C. perfringens.* In contrast, challenge of *C. perfringens* alone and in conjunction with *Eimeria* both promote proliferation of *Clostridium sensu stricto 1* and NE case. This indicates that the amount of commensal *C. perfringens* in the jejunum under *Eimeria* infection is not sufficient to reach the significant abundance of *Clostridium sensu stricto 1* or *C. perfringens*, subsequently promoting the occurrence of NE. Therefore, it is reasonable to suggest that the quantity of *Clostridium sensu stricto 1* or *C. perfringens in* jejunum is critical for the onset of proliferation. A recent study used commensal *C. perfringens*, the isolate from normal chicken, to challenge broiler and reproduce NE in conjunction with infection of *E. maxima* [64]. This result also highlighted that not the specific *C. perfringens* strain but the exogenous addition of *C. perfringens* played the key in achieving the consequence. Accordingly, the methodology to inhibit overgrowth of *Clostridium sensu stricto 1* or *C. perfringens* in small intestines will be the straightforward strategy to prevent NE.

Recent studies have been shown that cecal microbiota had a prominent role in feed efficiency [65] and received increasing attention in terms of diseases [66] and metabolism [67]. In this study, the result of PCoA and PCA demonstrated that microbial communities in the jejunum were different from those in the cecum. Side by side treatments of *C. perfringens* and *Eimeria* promoted microbial shifts with biological significance in the jejunum but minimal fluctuations in taxa abundance in the cecum. Comparatively, jejunal microbiota was more significant than cecal microbiota to address characteristic gut microbiota contributory to NE by means of metagenomeSeq and LEfSe analysis. The reason might be that cecal microbiota is demonstrated more diverse than other intestinal sections [68] and inhibits higher amounts of microbes (10^10^-10^11^ CFU/g) than those in the jejunum (10^8^-10^9^ CFU/g) [69]. Those may provide the buffer effect on microbial changes in cecal microbiota. Besides, preferential colonization of *C. perfringens* on mucosa of the small intestine [33] may also contribute to less amount of *C. perfringens* into cecum, hence adverse to elicit significant changes in cecal microbiota.

Medium-chain fatty acids (MCFAs) such as lauric acid are a family of saturated 6- to 12-carbon fatty acids from plants and documented beneficial effects on intestinal health and microbial growth inhibition [70–72]. The mechanism for their bactericidal activity is not fully understood. Relative studies showed that they could act as nonionic surfactants to become incorporated into the bacterial cell membrane, as well as diffuse through cell membranes and create pores, changing membrane permeability and leading to cell death [73–75]. In this work, lauric acid attracted interest due to its inexpensiveness and natural properties, including strong antibacterial effects against *C. perfringens* and no inhibitory effect on *Eimeria* infection [76]. Based on Timbermont’s study, lauric acid was most effective in inhibiting the growth of *C. perfringens* strain *in vitro*. Given a supplementary dose of 0.4 kg/ton in feed caused a significant decrease in NE incidence (from 50% down to 25%) compared with the infected, untreated control group [29]. This study followed the dose and used experimental grade product of lauric acid to evaluate the effects on NE incidence and intestinal microbiota. However, the addition of lauric acid did not reduce the incidence of NE. For intestinal microbiota, lauric acid neither exerted the inhibitory effect against proliferation of *C. perfringens* nor elevated the level of beneficial bacteria, such as *Lactobacillus* and *Bifidobacterium*. But, the relative abundance of *Escherichia Shigella* was decreased without affecting the incidence. Since lauric acid has different grade of products, such as experimental or food grade, the contradictory result may attribute to the influence of different formula on the absorptive efficiency of this compound. MCFAs are hydrophobic and partly absorbed through the stomach mucosa. Hence, their triacylglycerols are considered as a desirable formula for feed additive because they can be absorbed intact into intestinal epithelial enterocytes via this form [77].

In summary, significant overgrowth of *Clostridium sensu stricto 1* in jejunum was recognized as the major microbiota contributory to NE. In addition to *C. perfringens*, other member within *Clostridium sensu stricto 1* was also found to participate in disease development. The decrement of *Lactobacillus* following the NE severity indicated that lactobacilli also participate in the progress of disease. These taxa showed counteractive effects in their functions as well as in the bacterial abundance, attempting to maintain the homeostasis of jejunal microbiota in chickens. Therefore, manipulations to inhibit multiplication of *Clostridium sensu stricto 1* and *C. perfringens* and to rehabilitate the dominant *Lactobacillus* population in the jejunum should be the niche for developing effective strategies to prevent NE.

## Acknowledgments

The authors would like to thank Dr. John F. Prescott (University of Guelph, Ontario, Canada) for providing *netB* positive *C. perfringens* strain CP1. This work was supported by the USDA, National Institute of Food and Agriculture (CRIS Project Accession Number 1014508) and the College of Veterinary Medicine, Mississippi State University.

## Supporting information

**Figure S1**. Rarefaction curve by groups.

**Figure S2**. Comparison of genera abundance between jejunal and cecal microbiota by STAMP with Welch’s t-test.

**Figure S3**. Rank abundance curve by groups.

**Figure S4**. Comparison of genera (A) and species (B) abundance between AJ and EJ, BJ and EJ, and CJ and EJ groups by STAMP with Welch’s t-test.

